# A single intranasal dose of chimpanzee adenovirus-vectored vaccine confers sterilizing immunity against SARS-CoV-2 infection

**DOI:** 10.1101/2020.07.16.205088

**Authors:** Ahmed O. Hassan, Natasha M. Kafai, Igor P. Dmitriev, Julie M. Fox, Brittany Smith, Ian B. Harvey, Rita E. Chen, Emma S. Winkler, Alex W. Wessel, James Brett Case, Elena Kashentseva, Broc T. McCune, Adam L. Bailey, Haiyan Zhao, Laura A. VanBlargan, Yanan Dai, Meisheng Ma, Lucas J. Adams, Swathi Shrihari, Lisa E. Gralinski, Yixuan J. Hou, Alexandra Schaefer, Arthur S. Kim, Shamus P. Keeler, Daniela Weiskopf, Ralph Baric, Michael J. Holtzman, Daved H. Fremont, David T. Curiel, Michael S. Diamond

## Abstract

The Coronavirus Disease 2019 pandemic has made deployment of an effective vaccine a global health priority. We evaluated the protective activity of a chimpanzee adenovirus-vectored vaccine encoding a pre-fusion stabilized spike protein (ChAd-SARS-CoV-2-S) in challenge studies with Severe acute respiratory syndrome coronavirus 2 (SARS-CoV-2) and mice expressing the human angiotensin-converting enzyme 2 receptor. Intramuscular dosing of ChAd-SARS-CoV-2-S induces robust systemic humoral and cell-mediated immune responses and protects against lung infection, inflammation, and pathology but does not confer sterilizing immunity, as evidenced by detection of viral RNA and induction of anti-nucleoprotein antibodies after SARS-CoV-2 challenge. In contrast, a single intranasal dose of ChAd-SARS-CoV-2-S induces high levels of systemic and mucosal IgA and T cell responses, completely prevents SARS-CoV-2 infection in the upper and lower respiratory tracts, and likely confers sterilizing immunity in most animals. Intranasal administration of ChAd-SARS-CoV-2-S is a candidate for preventing SARS-CoV-2 infection and transmission, and curtailing pandemic spread.

## INTRODUCTION

Severe acute respiratory syndrome coronavirus 2 (SARS-CoV-2) is a positive-sense single-stranded RNA virus that was first isolated in late 2019 from patients with severe respiratory illness in Wuhan, China (Zhou et al., 2020b). As a betacoronavirus, SARS-CoV-2 is related to two other highly pathogenic respiratory viruses, SARS-CoV and Middle East respiratory syndrome coronavirus (MERS-CoV). SARS-CoV-2 infection results in a clinical syndrome, Coronavirus Disease 2019 (COVID-19) that can progress to respiratory failure (Guan et al., 2020) and also present with cardiac pathology, gastrointestinal disease, coagulopathy, and a hyperinflammatory syndrome (Cheung et al., 2020; Mao et al., 2020; Wichmann et al., 2020). The elderly, immunocompromised, and those with co-morbidities (*e*.*g*., obesity, diabetes, and hypertension) are at greatest risk of death from COVID-19 (Zhou et al., 2020a). Virtually all countries and territories have been affected with more than twelve million infections to date, hundreds of thousands of deaths, and a case-fatality rate estimated at ∼4%. Most cases are spread by direct human-to-human and droplet contact with substantial transmission in asymptomatic or mildly symptomatic individuals (Furukawa et al., 2020; Li et al., 2020). The extensive morbidity, mortality, and destabilizing socioeconomic consequences of COVID-19 highlight the urgent need for deployment of an effective SARS-CoV-2 vaccine to mitigate the severity of infection, curb transmission, end the pandemic, and prevent its return.

The SARS-CoV-2 RNA genome is approximately 30,000 nucleotides in length. The 5’ two-thirds encode nonstructural proteins that enable genome replication and viral RNA synthesis. The remaining one-third encode structural proteins such as spike (S), envelope, membrane, and nucleoprotein (NP) that form the spherical virion, and accessory proteins that regulate cellular responses. The S protein forms homotrimeric spikes on the virion and engages the cell-surface receptor angiotensin-converting enzyme 2 (ACE2) to promote coronavirus entry into human cells (de Wit et al., 2016; Letko et al., 2020). The SARS-CoV and SARS-CoV-2 S proteins are cleaved sequentially during the entry process to yield S1 and S2 fragments, followed by further processing of S2 to yield a smaller S2’ protein (Hoffmann et al., 2020). The S1 protein includes the receptor binding domain (RBD) and the S2 protein promotes membrane fusion. The structure of a soluble, stabilized prefusion form of the SARS-CoV-2 S protein was solved by cryo-electron microscopy, revealing considerable similarity to the SARS-CoV S protein (Wrapp et al., 2020b). This form of the S protein is recognized by potently neutralizing monoclonal antibodies (Cao et al., 2020; Pinto et al., 2020; Zost et al., 2020) and could serve as a promising vaccine target.

Release of the SARS-CoV-2 genome sequence prompted academic, government, and industry groups to immediately begin developing vaccine candidates that principally targeted the viral S protein (Burton and Walker, 2020). Multiple platforms have been developed to deliver the SARS-CoV-2 S protein including DNA plasmid, lipid nanoparticle encapsulated mRNA, inactivated virion, and viral-vectored vaccines (Graham, 2020). Several vaccines have entered clinical trials to evaluate safety, and some have advanced to trials assessing immunogenicity and efficacy. Because of the urgency of the pandemic, most vaccines advanced to human testing without substantive efficacy data in animals (Diamond and Pierson, 2020). This circumstance occurred in part because vaccine design and development has outpaced the generation of accessible pre-clinical disease models of SARS-CoV-2 infection and pathogenesis.

Adenovirus (Ad)-based vaccines against betacoronaviruses have been evaluated previously. A single dose of a chimpanzee Ad-vectored vaccine encoding the full-length S protein of MERS-CoV protected human dipeptidyl peptidase 4 (hDPP4) transgenic mice from infection (Munster et al., 2017), reduced virus shedding and enhanced survival in camels (Alharbi et al., 2019), and was safe and immunogenic in humans in a phase 1 clinical trial (Folegatti et al., 2020). A human Ad-based vaccine expressing a MERS S1-CD40L fusion protein also was protective in transgenic hDPP4 mice (Hashem et al., 2019). An Ad-based SARS-CoV vaccine expressing a full-length S protein prevented pneumonia in ferrets after challenge and was highly immunogenic in rhesus macaques (Kobinger et al., 2007). A chimpanzee Ad vector (Y25, a simian Ad-23 (Dicks et al., 2012)) encoding the wild-type SARS-CoV-2 S protein (ChAdOx1 nCoV-19) is currently under evaluation in humans as a single intramuscular injection (NCT04324606). Preliminary pre-print analysis suggests this vaccine protects against lung infection and pneumonia but not against upper respiratory tract infection and nasal virus shedding (https://doi.org/10.1101/2020.05.13.093195).

Here, we develop and evaluate a different chimpanzee Ad (simian AdV-36)-based SARS-CoV-2 vaccine (ChAd-SARS-CoV-2-S) encoding a pre-fusion stabilized spike (S) protein after introducing two proline substitutions in the S2 subunit (Pallesen et al., 2017). Intramuscular administration of ChAd-SARS-CoV-2-S induced robust systemic humoral and cell-mediated immune responses against the S protein. One or two vaccine doses protected against lung infection, inflammation, and pathology after SARS-CoV-2 challenge of mice that transiently express the human ACE2 (hACE2) receptor. Despite the induction of high levels of neutralizing antibody in serum, neither dosing regimen completely protected against SARS-CoV-2 infection, as substantial levels of viral RNA were still detected in the lung. In comparison, when we administered a single dose of ChAd-SARS-CoV-2-S by the intranasal route, we detected high levels of neutralizing antibody and anti-SARS-CoV-2 IgA, and complete protection against infection in both the upper and lower respiratory tracts. Moreover, and in contrast to the control ChAd vaccine, 8 days after SARS-CoV-2 challenge, serum antibody responses to the SARS-CoV-2 NP protein were absent in animals immunized with ChAd-SARS-CoV-2-S via intranasal delivery. Thus, ChAd-SARS-CoV-2-S has the potential to confer sterilizing immunity at the site of inoculation, which should prevent both virus-induced disease and transmission.

## RESULTS

### Chimpanzee Ad-vectored vaccine induces robust antibody responses against anti-SARS-CoV-2

We constructed two replication-incompetent ChAd vectors based on a simian Ad-36 virus. The ChAd-SARS-CoV-2-S vector encodes the full-length sequence of SARS-CoV-2 S protein as a transgene including the ectodomain, transmembrane domain, and cytoplasmic domain (GenBank: QJQ84760.1) and is stabilized in pre-fusion form by two proline substitutions at residues K986 and V987 (Pallesen et al., 2017; Wrapp et al., 2020a). The ChAd-control has no transgene. The S protein transgene is controlled transcriptionally by a cytomegalovirus promoter. To make the vector replication-incompetent and enhance packaging capacity, we replaced the E1A/B genes and introduced a deletion in the E3B gene, respectively (**Fig 1A**). To confirm that the S protein was expressed and antigenically intact, we transduced 293T cells and confirmed binding of a panel of 22 neutralizing monoclonal antibodies against the S protein by flow cytometry (**Fig 1B**).

**Figure 1.**
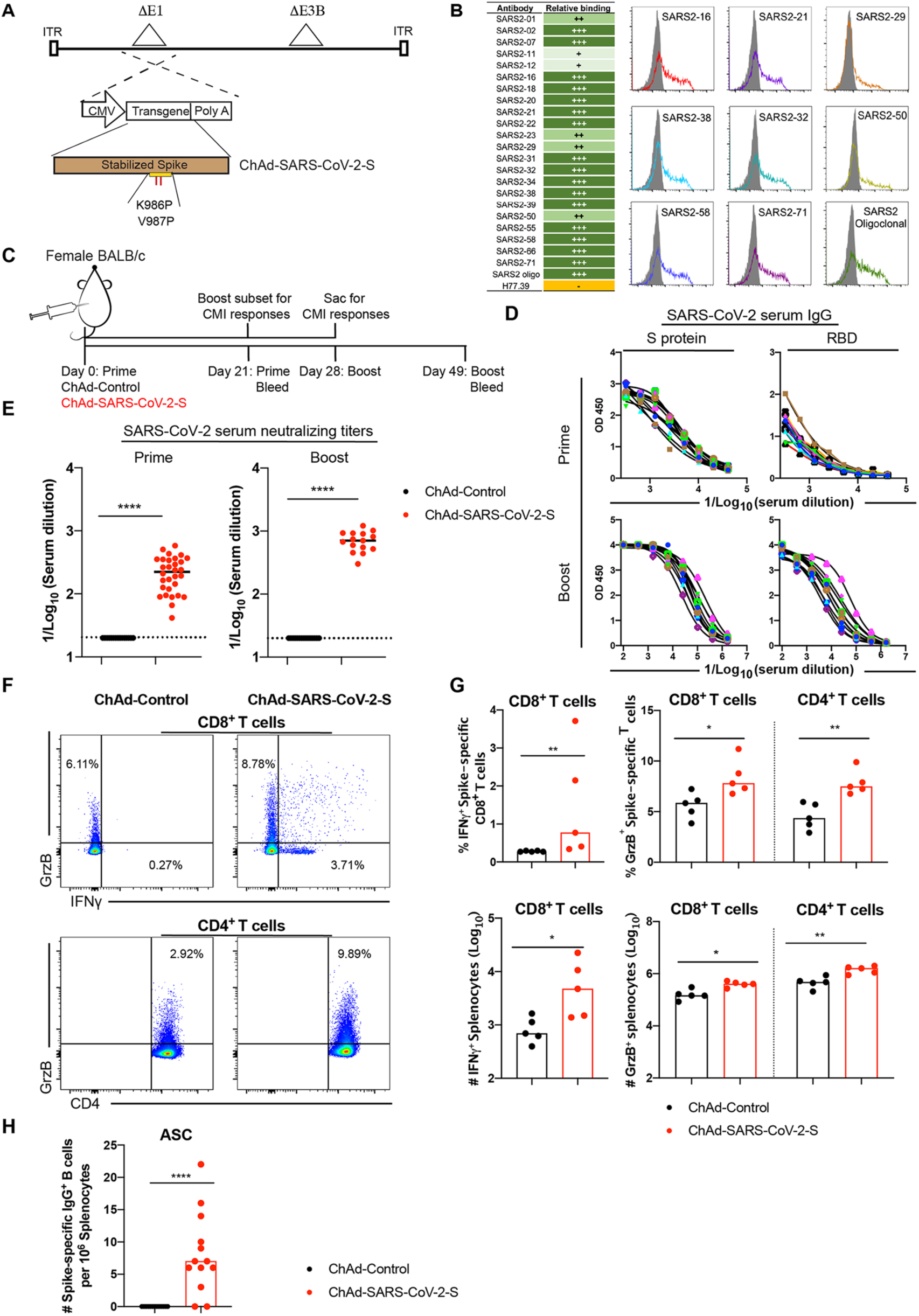
Immunogenicity of ChAd-SARS-CoV-2-S. **A.** Diagram of transgene cassettes: ChAd-control has no transgene insert; ChAd-SARS-CoV-2-S encodes for SARS-CoV-2 S protein with the two indicated proline mutations. **B**. Binding of ChAd-SARS-CoV-2-S transduced 293 cells with anti-S mAbs. (*Left*) Summary: +, ++, +++, - indicate < 25%, 25-50%, > 50%, and no binding, respectively. (*Right*) Representative flow cytometry histograms of two experiments. **C-E**. Four-week old female BALB/c mice were immunized via intramuscular route with ChAd-control or ChAd-SARS-CoV-2-S and boosted four weeks later. Antibody responses in sera of immunized mice at day 21 after priming or boosting were evaluated. An ELISA measured anti-S and RBD IgG levels (**D**), and an FRNT determined neutralization activity (**E**). Data are pooled from two experiments (n = 15 to 30; Mann-Whitney test: ****, *P* < 0.0001). **F-G**. Cell-mediated responses were analyzed at day 7 post-booster immunization after re-stimulation with an S protein peptide pool (**Table S1**). Splenocytes were assayed for IFNγ and granzyme B expression in CD8^+^ T cells and granzyme B only in CD4^+^ T cells by flow cytometry (**F**). A summary of frequencies and numbers of positive cell populations is shown in **G** (n = 5; Mann-Whitney test: *, *P* < 0.05; **, *P* < 0.01; ***, *P* < 0.001). Bars indicate median values, and dotted lines are the limit of detection (LOD) of the assays. **H**. Spleens were harvested at 7 days post-boost, and SARS-CoV-2 spike-specific IgG+ antibody-secreting cells (ASC) frequency was measured by ELISPOT (Mann-Whitney test: ****, *P* < 0.0001). Bars and columns show median values, and dotted lines indicate the limit of detection (LOD) of the assays.

To assess the immunogenicity of ChAd-SARS-CoV-2-S, groups of 4-week-old BALB/c mice were immunized by intramuscular inoculation with 10^10^ virus particles of ChAd-SARS-CoV-2-S or ChAd-control. Some mice received a booster dose four weeks later. Serum samples were collected 21 days after primary or booster immunization (**Fig 1C**), and IgG responses against purified S and RBD proteins were evaluated by ELISA. Whereas ChAd-SARS-CoV-2-S induced high levels of S- and RBD-specific IgG, low, if any levels were detected in the ChAd-control-immunized mice (**Fig 1D and S1A**). Serum samples were assayed *in vitro* for neutralization of infectious SARS-CoV-2 using a focus-reduction neutralization test (FRNT) (Case et al., 2020). As expected, serum from ChAd-control-immunized mice did not inhibit SARS-CoV-2 infection after primary immunization or boosting. In contrast, serum from ChAd-SARS-CoV-2-S vaccinated animals strongly neutralized SARS-CoV-2 infection, and boosting enhanced this inhibitory activity (**Fig 1E and S1B-C**).

### Vaccine-induced memory CD8^+^ T cell and antigen specific B cell responses

Because optimal vaccine immunity often is comprised of both humoral and cellular responses (Slifka and Amanna, 2014), we measured the levels of SARS-CoV-2-specific CD4^+^ and CD8^+^ T cells after vaccination. Four-week old BALB/c mice were immunized with ChAd-SARS-CoV-2-S or ChAd-control and boosted three weeks later. To assess the vaccine-induced SARS-CoV-2-specific CD4^+^ and CD8^+^ T cell responses, splenocytes were harvested one week after boosting and stimulated *ex vivo* with a pool of 253 overlapping 15-mer S peptides (**Table S1**). Subsequently, quantification of intracellular IFNγ and granzyme B expression was determined by flow cytometry. After peptide re-stimulation *ex vivo*, splenic CD8^+^ T cells expressed IFNγ and both splenic CD4^+^ and CD8^+^ T cells expressed granzyme B in mice immunized with ChAd-SARS-CoV-2-S but not the ChAd-control vector (**Fig 1F-G and S2**). To assess the antigen-specific B cell responses, splenocytes were harvested and subjected to an ELISPOT analysis with S protein. The ChAd-SARS-CoV-2-S vaccine induced S protein-specific IgG antibody-secreting cells in the spleen whereas the control vaccine did not (**Fig 1H**).

### Intramuscular immunization with ChAd-SARS-CoV-2-S vaccine protects against SARS-CoV-2 infection in the lung

We tested the protective activity of the ChAd vaccine in a recently developed SARS-CoV-2 infection model wherein BALB/c mice express hACE2 in the lung after intranasal delivery of a vectored human Ad (Hu-Ad5-hACE2) (Hassan et al., 2020). Endogenous mouse ACE2 does not support viral entry (Letko et al., 2020), and this system enables productive SARS-CoV-2 infection in the mouse lung. Four-week-old BALB/c mice first were immunized via an intramuscular route with ChAd-control or ChAd-SARS-CoV-2-S vaccines. Approximately thirty days later, mice were administered 10^8^ plaque-forming units (PFU) of Hu-Ad5-hACE2 and anti-Ifnar1 monoclonal antibody (mAb) via intranasal and intraperitoneal routes, respectively. We also administered a single dose of anti-Ifnar1 mAb to enhance lung pathogenesis in this model (Hassan et al., 2020). We confirmed the absence of cross-immunity between the ChAd and the Hu-Ad5 vector. Serum from ChAd-immunized mice did not neutralize Hu-Ad5 infection (**Fig S3A-B**).

Five days after Hu-Ad5-hACE2 transduction, mice were challenged via intranasal route with 4 × 10^5^ focus-forming units (FFU) of SARS-CoV-2 (**Fig 2A**). At 4 days post-infection (dpi), the peak of viral burden in this model (Hassan et al., 2020), mice were euthanized, and lungs, spleen, and heart were harvested for viral burden and cytokine analysis. Notably, there was no detectable infectious virus in the lungs of mice immunized with ChAd-SARS-CoV-2-S as determined by plaque assay, whereas high levels were present in mice vaccinated with ChAd-control (**Fig 2B**). Consistent with this result, we observed reduced viral RNA levels in the lung, heart, and spleen of ChAd-SARS-CoV-2-S vaccinated animals compared to mice receiving the ChAd-control vector (**Fig 2C**). *In situ* hybridization staining for viral RNA in lungs harvested at 4 dpi revealed a substantial decrease of SARS-CoV-2 RNA in pneumocytes of animals immunized with ChAd-SARS-CoV-2-S compared to the ChAd-control (**Fig 2D**). A subset of immunized animals was euthanized at 8 dpi, and tissues were harvested for evaluation. At this time point, viral RNA levels again were lower or absent in the lung and spleen of ChAd-SARS-CoV-2-S immunized mice compared to the control ChAd vector (**Fig 2C**). Collectively, these data indicate that a single intramuscular immunization with ChAd-SARS-CoV-2-S results in markedly reduced, but not abrogated, SARS-CoV-2 infection in the lungs of challenged mice.

**Figure 2.**
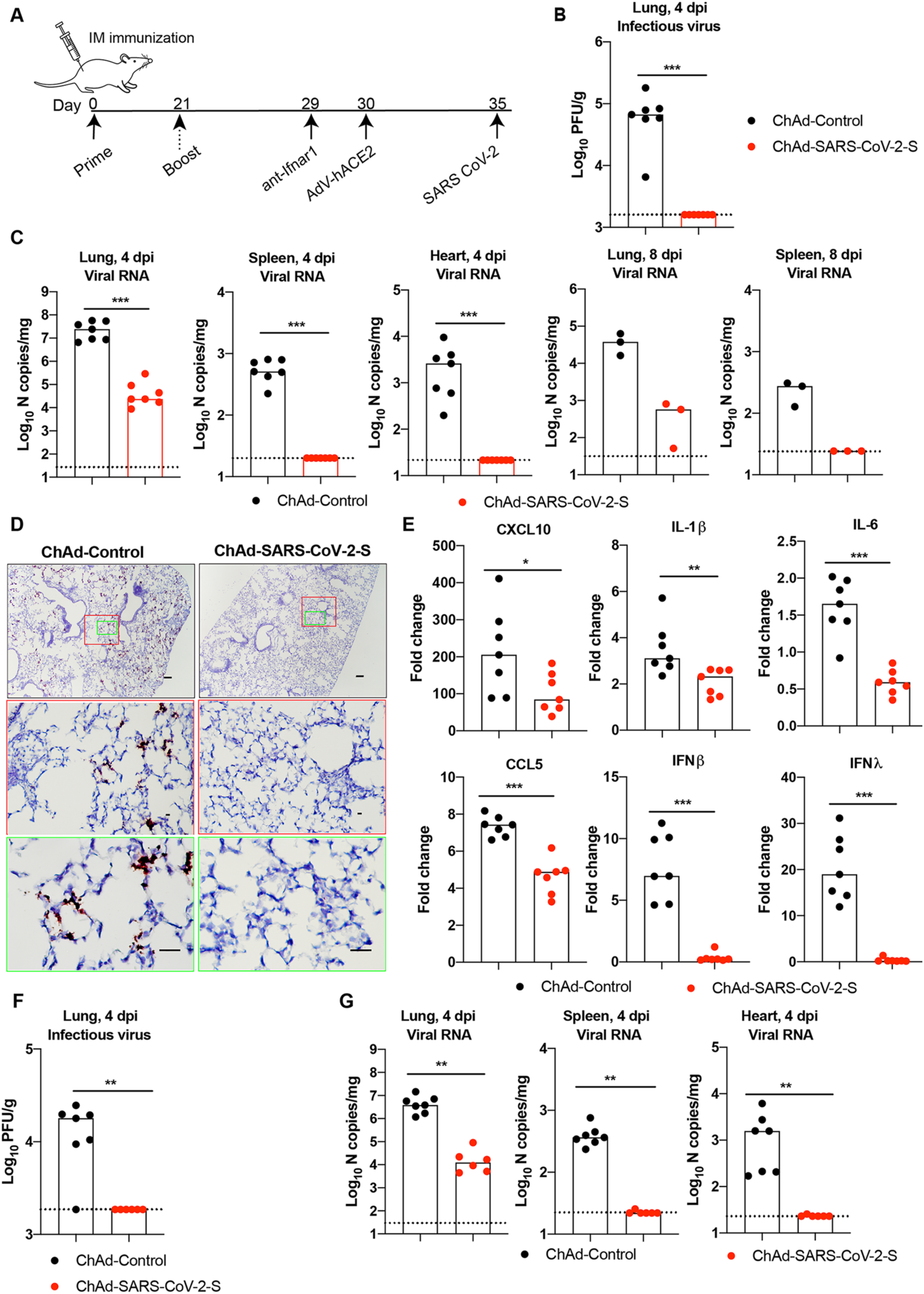
Protective efficacy of intramuscularly delivered ChAd-SARS-CoV-2-S against SARS-CoV-2 infection. **A**. Scheme of vaccination and challenge. Four-week-old BALB/c female mice were immunized ChAd-control or ChAd-SARS-CoV-2-S. Some mice received a booster dose of the homologous vaccine. On day 35 post-immunization, mice were challenged with SARS-CoV-2 as follows: animals were treated with anti-Ifnar1 mAb and transduced with Hu-AdV5-hACE2 via an intranasal route one day later. Five days later, mice were challenged with 4 × 10^5^ focus-forming units (FFU) of SARS-CoV-2 via the intranasal route. **B-C**. Tissues were harvested at 4 and 8 dpi for analysis. Infectious virus in the lung was measured by plaque assay (**B**) and viral RNA levels were measured in the lung, spleen and heart at 4 and 8 dpi by RT-qPCR (**C**) (n = 3-7, Mann-Whitney test: *** *P* < 0.001). **D**. Viral RNA *in situ* hybridization using SARS-CoV-2 probe (brown color) in the lungs harvested at 4 dpi. Images show low-(top; scale bars, 100 μm) and medium-(middle; scale bars, 100 μm) power magnification with a high-power magnification inset (representative images from n = 3 per group). **E**. Fold change in gene expression of indicated cytokines and chemokines from lung homogenates at 4 dpi was determined by RT-qPCR after normalization to *Gapdh* levels and comparison with naïve unvaccinated, unchallenged controls (n = 7; Mann-Whitney test: ***, *P* < 0.001). **F-G**. Mice that received a prime-boost immunization were challenged on day 35 post-booster immunization. Tissues were collected at 4 dpi for analysis. Infectious virus in the lung was determined by plaque assay (**F**) and viral RNA was measured in the lung, spleen and heart using RT-qPCR (**G**) (n = 6-7; Mann-Whitney test: **, *P* < 0.01). (**B-C** and **E-G**) Columns show median values, and dotted lines indicate the LOD of the assays.

We next assessed the effect of the vaccine on lung inflammation and disease. Several proinflammatory cytokines and chemokine mRNA levels were lower in the lung tissues of animals immunized with ChAd-SARS-CoV-2-S compared to ChAd-control including *CXCL10, IL1β, IL-6, CCL5, IFNβ*, and *IFNλ* (**Fig 2E**). Moreover, mice immunized with ChAd-control vaccine and challenged with SARS-CoV-2 showed evidence of viral pneumonia characterized by immune cell accumulation in perivascular and alveolar locations, vascular congestion, and interstitial edema. In contrast, animals immunized with ChAd-SARS-CoV-2-S showed a marked attenuation of the inflammatory response in the lung that develops in the ChAd-control-immunized mice (**Fig 3**). Thus, immunization with Ch-Ad-SARS-CoV-2 decreases both viral infection and the consequent lung inflammation and injury associated with SARS-CoV-2 infection.

**Figure 3.**
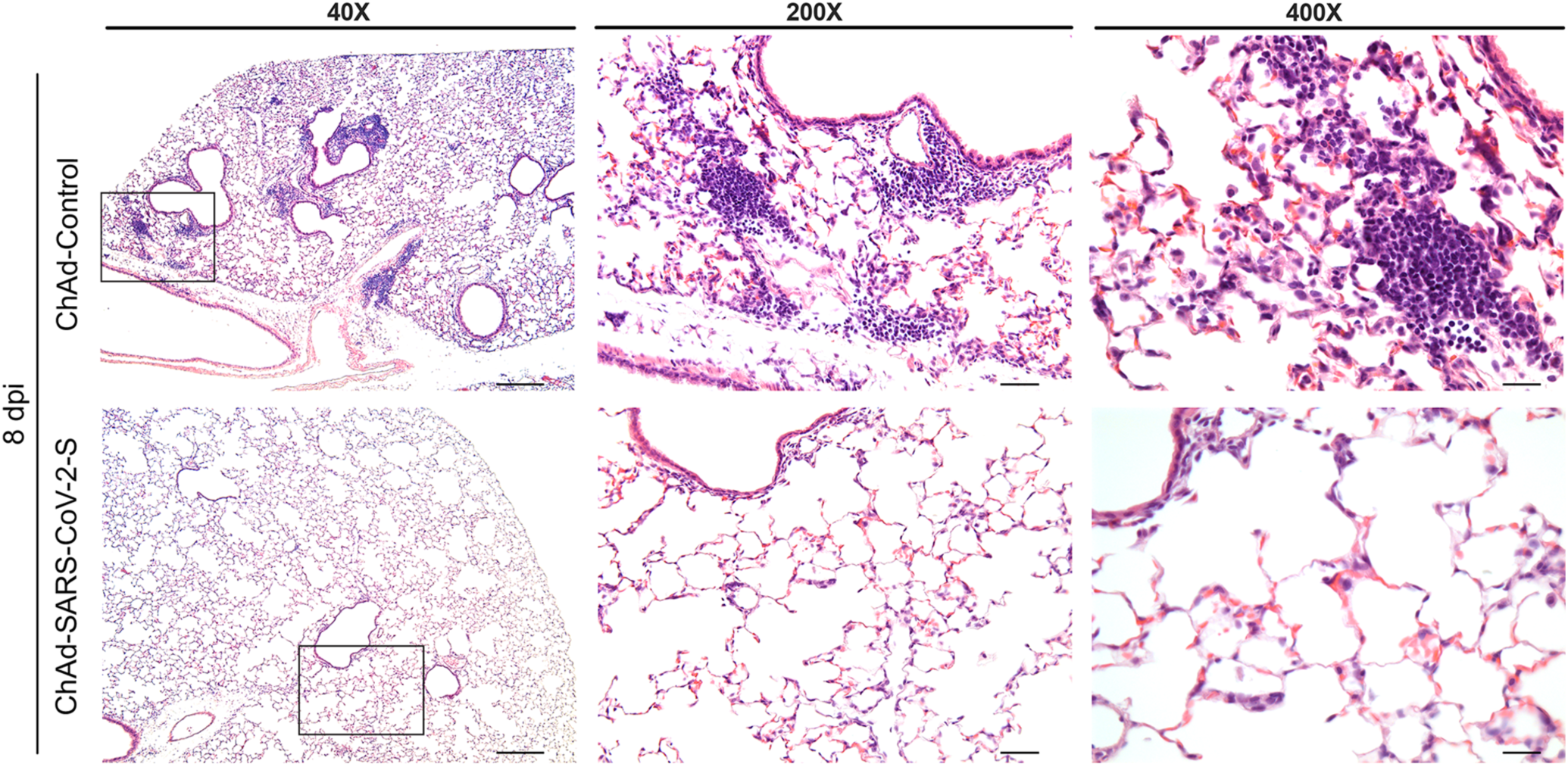
Single-dose intramuscular vaccination with ChAd-SARS-CoV-2-S protects mice against SARS-CoV-2-induced inflammation in the lung. Four-week old female BALB/c mice were immunized with ChAd-control and ChAd-SARS-CoV-2-S and challenged following the scheme described in **Figure 2**. Lungs were harvested at 8 dpi. Sections were stained with hematoxylin and eosin and imaged at 40x (left; scale bar, 250 μm), 200x (middle; scale, 50 μm), and 400x (right; scale bar, 25 μm) magnifications. Each image is representative of a group of 3 mice.

We then assessed for improved protection using a prime-boost vaccine regimen. BALB/c mice were immunized via an intramuscular route with ChAd-control or ChAd-SARS-CoV-2-S and received a homologous booster dose four weeks later. At day 29 post-boost, mice were treated with a single dose of anti-Ifnar1 antibody followed by Hu-Ad5-hACE2 and then challenged with SARS-CoV-2 five days later. As expected, the prime-boost regimen protected against SARS-CoV-2 challenge with no infectious virus detected in the lungs (**Fig 2F**). Although marked reductions of viral RNA in the lung, spleen, and heart were detected at 4 dpi, residual levels of viral RNA still were present suggesting protection was not complete, even after boosting (**Fig 2G**).

### A single intranasal immunization with ChAd-SARS-CoV-2-S induces sterilizing immunity against SARS-CoV-2

Mucosal immunization through the nasopharyngeal route can elicit local immune responses including secretory IgA antibodies that confer protection at or near the site of inoculation of respiratory pathogens (Neutra and Kozlowski, 2006). To assess the immunogenicity and protective efficacy of mucosal vaccination, five-week old BALB/c mice were inoculated via intranasal route with 10^10^ viral particles of ChAd-control or ChAd-SARS-CoV-2-S (**Fig 4A**). Serum samples were collected at four weeks post-immunization to evaluate humoral immune responses. Intranasal immunization of ChAd-SARS-CoV-2-S but not ChAd-control induced high levels of S- and RBD-specific IgG and IgA (**Fig 4B-C**) and SARS-CoV-2 neutralizing antibodies (geometric mean titer of 1/1,574) in serum (**Fig 4D and S4A**). Serum antibodies from mice immunized with ChAd-SARS-CoV-2-S equivalently neutralized a recombinant, luciferase-expressing variant of SARS-CoV-2 encoding a D614G mutation in the S protein (**Fig S4B**); this finding is important, because many circulating viruses contain this substitution, which is associated with greater infectivity in cell culture (https://doi.org/10.1101/2020.06.12.148726). We also assessed the SARS-CoV-2-specific antibody responses in bronchoalveolar lavage (BAL) fluid of immunized mice. BAL fluid from ChAd-SARS-CoV-2-S but not ChAd-control vaccinated mice showed high levels of S- and RBD-specific IgG and IgA antibodies (**Fig 4E-F**) including those with neutralizing activity (**Fig 4G and S4C**).

**Figure 4.**
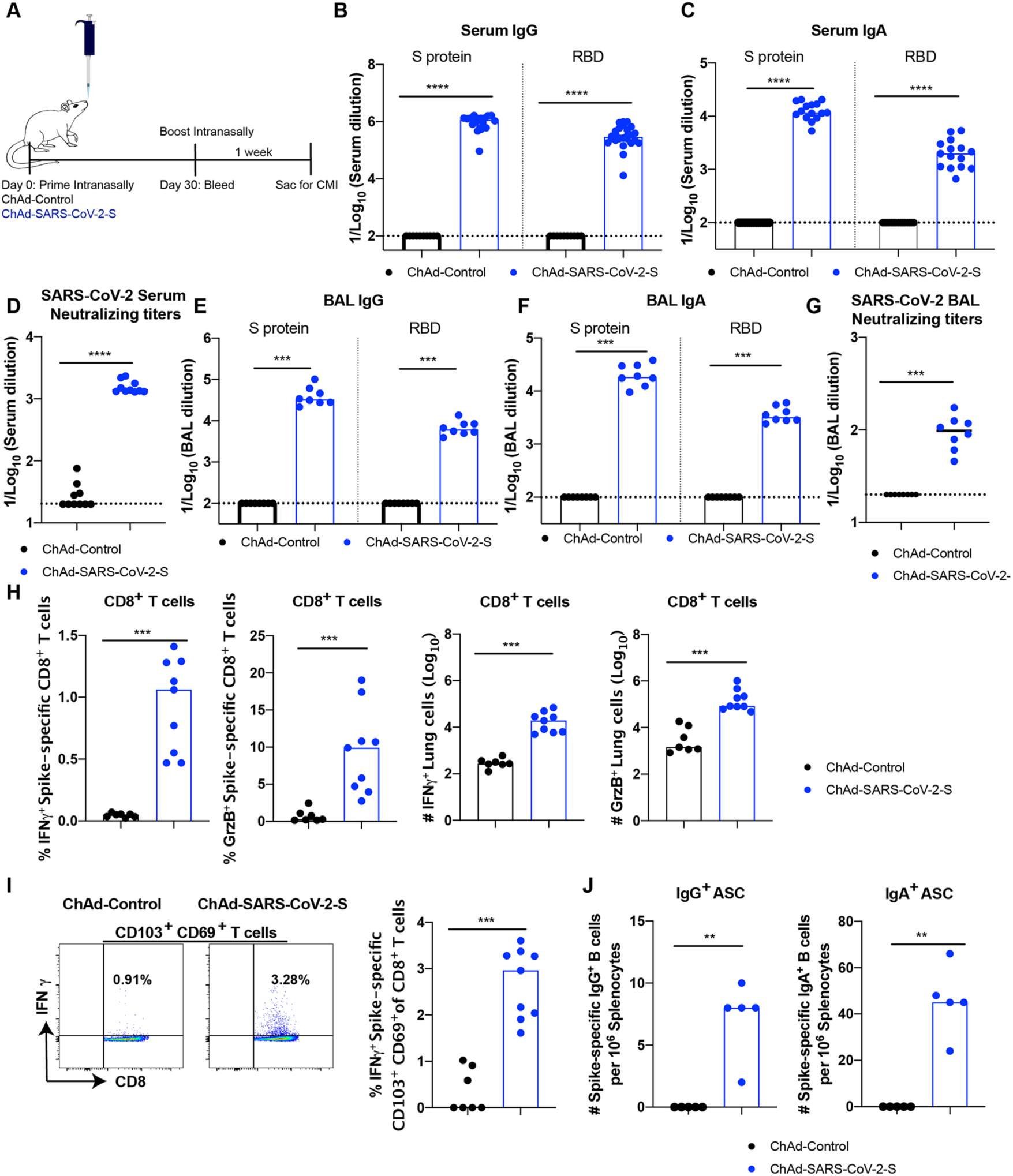
Immune responses after Intranasal immunization of ChAd-SARS-CoV-2-S. **A**. Scheme of experiments. Five-week-old BALB/c female mice were immunized with ChAd-control or ChAd-SARS-CoV-2-S via an intranasal route. (**B-D**) Antibody responses in sera of immunized mice at one month after priming were evaluated. An ELISA measured SARS-CoV-2 S-and RBD-specific IgG (**B**) and IgA levels (**C**), and a FRNT determined neutralization activity (**D**). Data are pooled from two experiments with n = 10-25 mice per group (Mann-Whitney test: ****, *P* < 0.0001). **E-J**. Mice that received a booster dose were sacrificed one week later to evaluate mucosal and cell-mediated immune responses. SARS-CoV-2 S-and RBD-specific IgG (**E**) and IgA levels (**F**) in BAL fluid were determined by ELISA. Neutralizing activity of BAL fluid against SARS-CoV-2 was measured by FRNT (**G**). CD8^+^ T cells in the lung were assayed for IFNγ and granzyme B expression by flow cytometry after re-stimulation with an S protein peptide pool (**H**). CD8^+^ T cells in the lung also were phenotyped for expression of CD103 and CD69 (**I**). SARS-CoV-2 spike-specific IgG^+^ and IgA^+^ antibody-secreting cells (ASC) frequency in the spleen harvested one week post-boost was measured by ELISPOT (**J**). Data for mucosal and cell-mediated responses are pooled from two experiments (**E-I**: n = 7-9 per group; Mann-Whitney test: ***, *P* < 0.001); **J**: n = 5 per group; Mann-Whitney test: **, *P* < 0.01; ***, *P* < 0.001). (**B-J**) Bars and columns show median values, and dotted lines indicate the LOD of the assays.

To assess T cell responses activated via mucosal immunization, mice were vaccinated via intranasal route with either ChAd-SARS-CoV-2-S or ChAd-control and boosted similarly four weeks later. Lungs were harvested one-week post-boosting, and T cells were analyzed by flow cytometry. Re-stimulation *ex vivo* with a pool of S peptides resulted in a marked increase in IFNγ and granzyme B producing CD8^+^ T cells in the lungs of mice that received the ChAd-SARS-CoV-2-S vaccine (**Fig 4H**). Specifically, a population of IFNγ-secreting, antigen-specific CD103^+^CD69^+^CD8^+^ T cells in the lung was identified (**Fig 4I**) which is phenotypically consistent with vaccine-induced resident memory T cells (Takamura, 2017). In the spleen, we detected antibody-secreting plasma cells producing IgA or IgG against the S protein after intranasal immunization with ChAd-SARS-CoV-2-S (**Fig 4J**). Of note, we observed an ∼-5-fold higher frequency of B cells secreting anti-S IgA than IgG.

We evaluated the protective efficacy of the ChAd vaccine after single-dose intranasal immunization. At day 30 post-vaccination, mice were administered 10^8^ PFU of Hu-Ad5-hACE2 and anti-Ifnar1 mAb as described above. Five days later, mice were challenged with 4 × 10^5^ FFU of SARS-CoV-2. At 4 and 8 dpi, lungs, spleen, heart, nasal turbinates, and nasal washes were harvested and assessed for viral burden. Intranasal delivery of the ChAd-SARS-CoV-2-S vaccine demonstrated remarkable protective efficacy as judged by an absence of infectious virus in the lungs (**Fig 5A**) and almost no measurable viral RNA in the lung, spleen, heart, nasal turbinates, or nasal washes (**Fig 5B**). The very low viral RNA levels in the lung and nasal turbinates at 4 dpi likely reflects the input, non-replicating virus, as similar levels were measured at this time point in C57BL/6 mice lacking hACE2 receptor expression (Hassan et al., 2020). Cytokine and chemokine mRNA levels in lung homogenates also were substantially lower in mice immunized with the ChAd-SARS-CoV-2-S than the ChAd-control vaccine (**Fig 5C**), with residual expression likely due to the human Ad vector hACE2 delivery system. Finally, histopathological analysis of lung tissues from animals vaccinated with ChAd-SARS-CoV-2-S by intranasal route and challenged with SARS-CoV-2 showed minimal, if any, perivascular and alveolar infiltrates at 8 dpi compared to the extensive inflammation observed in ChAd-control vaccinated animals (**Fig 5D**).

**Figure 5.**
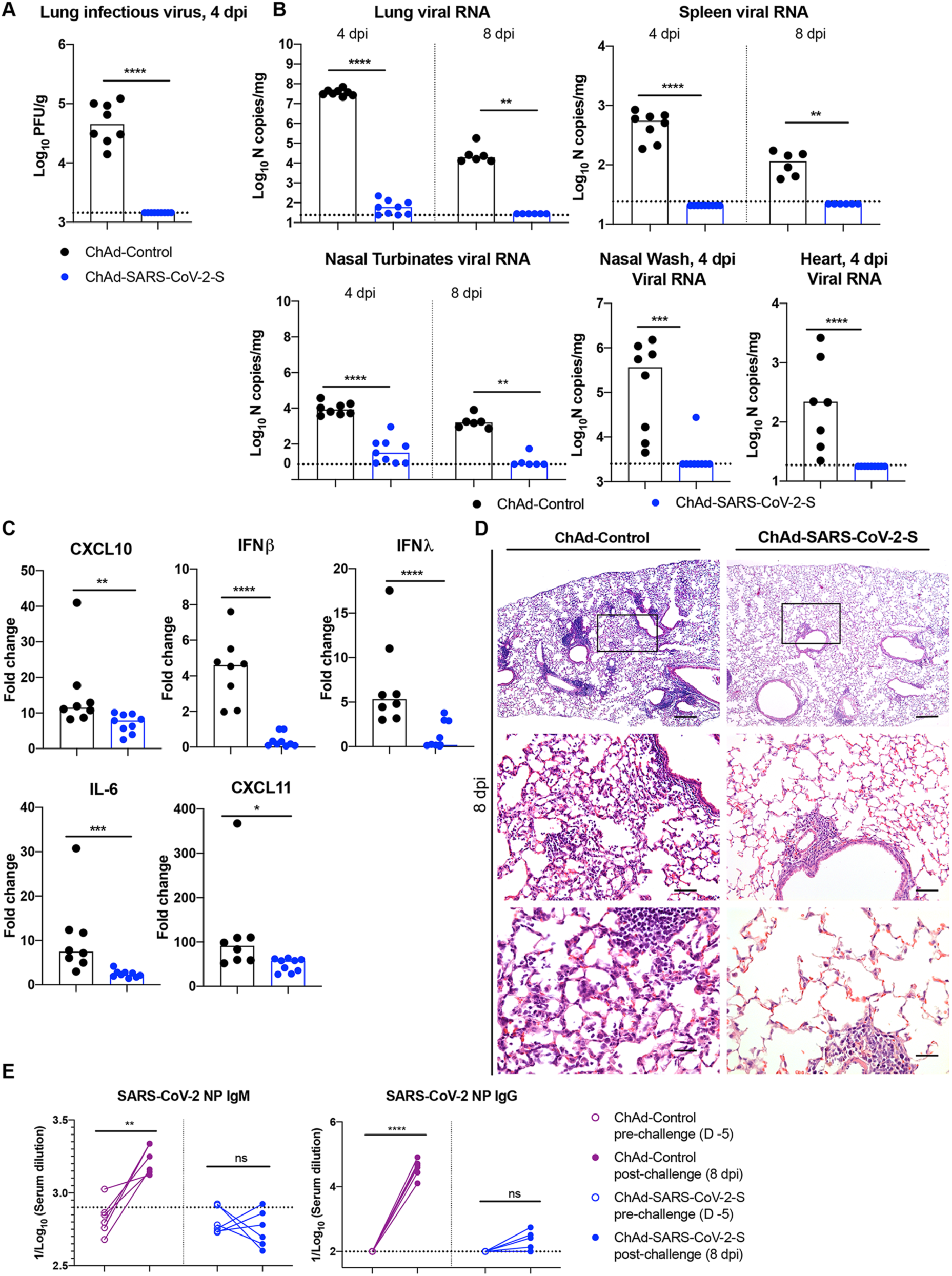
Single-dose intranasal immunization with ChAd-SARS-CoV-2-S protects against SARS-CoV-2 infection. Five-week-old BALB/c female mice were immunized with ChAd-control or ChAd-SARS-CoV-2-S via an intranasal route. On day 35 post-immunization, mice were challenged as follows: animals were treated with anti-Ifnar1 mAb and transduced with Hu-AdV5-hACE2 via the intranasal route one day later. Five days later, mice were challenged intranasally with 4 × 10^5^ FFU of SARS-CoV-2. (**A-C**) Tissues and nasal washes were collected at 4 and 8 dpi for analysis. Infectious virus in the lung was measured by plaque assay (**A**). Viral RNA levels in the lung, spleen, heart, nasal turbinates, and nasal washes were measured at 4 and 8 dpi by RT-qPCR (**B**). Fold change in gene expression of indicated cytokines and chemokines was determined by RT-qPCR, normalized to *Gapdh*, and compared to naïve controls in lung homogenates at 4 dpi (**C**) (2 experiments, n = 6-9; median values are shown: *, *P* < 0.05, ** *P* < 0.01, *** *P* < 0.001, **** *P* < 0.0001; Mann-Whitney test). Columns show median values, and dotted lines indicate the LOD of the assays. **D**. Lungs were harvested at 8 dpi. Sections were stained with hematoxylin and eosin and imaged at 40x (left; scale bar, 250 μm), 200x (middle; scale, 50 μm), and 400x (right; scale bar, 25 μm) magnifications. Each image is representative of a group of 3 mice. **E**. An ELISA measured anti-SARS-CoV-2 NP IgM (*left*) and IgG (*right*) antibody responses in paired sera obtained 5 days before and 8 days after SARS-CoV-2 challenge of ChAd-control or ChAd-SARS-CoV-2-S mice vaccinated by an intranasal route (n = 6: ns; not significant; ** *P* < 0.01, **** *P* < 0.0001; paired t test). Dotted lines represent the mean IgM and IgG titers from naïve sera (n = 6).

To determine if sterilizing immunity was achieved with intranasal delivery of ChAd-SARS-CoV-2-S, we measured anti-NP antibodies at 8 dpi and compared them to responses from 5 days before SARS-CoV-2 infection. Because the NP gene is absent from the vaccine vector, induction of humoral immune responses against NP after SARS-CoV-2 exposure (Ni et al., 2020) suggests viral protein translation and active infection. After SARS-CoV-2 challenge, anti-NP antibody responses were detected in ChAd-control mice vaccinated by an intranasal route or ChAd-control and ChAd-SARS-CoV-2-S mice vaccinated by an intramuscular route (**Fig 5E and S1D**). Remarkably, none of the mice immunized with ChAd-SARS-CoV-2-S via an intranasal route showed significant increases in anti-NP antibody responses after SARS-CoV-2 infection. Combined with our virological analyses, these data suggest that a single intranasal immunization of ChAd-SARS-CoV-2-S induces robust and likely sterilizing mucosal immunity, which prevents SARS-CoV-2 infection in the upper and lower respiratory tracts of mice expressing the hACE2 receptor.

## DISCUSSION

In this study, we evaluated intramuscular and intranasal administration of a replication-incompetent ChAd vector as a vaccine platforms for SARS-CoV-2. Single dose immunization of a stabilized S protein-based vaccine via an intramuscular route induced S- and RBD-specific binding as well as neutralizing antibodies. Vaccination with one or two doses protected mice expressing human ACE2 against SARS-CoV-2 challenge, as evidenced by an absence of infectious virus in the lungs and substantially reduced viral RNA levels in lungs and other organs. Mice immunized with ChAd-SARS-CoV-2-S also showed markedly reduced if not absent lung pathology, lung inflammation, and evidence of pneumonia compared to the control ChAd vaccine. However, intramuscular vaccination of ChAd-SARS-CoV-2-S did not confer sterilizing immunity, as evidenced by detectable viral RNA levels in several tissues including the lung and induction of anti-NP antibody responses. Mice immunized with a single dose of the ChAd-SARS-CoV-2-S via an intranasal route also were protected against SARS-CoV-2 challenge. Intranasal vaccination, however, generated robust IgA and neutralizing antibody responses that protected against both upper and lower respiratory tract SARS-CoV-2 infection and inhibited infection of both wild-type and D614G variant viruses. The very low viral RNA in upper airway tissues and absence of serological response to NP in the context of challenge strongly suggests that most animals receiving a single intranasal dose of ChAd-SARS-CoV-2-S achieve sterilizing immunity.

Although several vaccine candidates (*e*.*g*., lipid-encapsulated mRNA, DNA, inactivated, and viral-vectored) rapidly advanced to human clinical trials in an expedited effort to control the pandemic (Amanat and Krammer, 2020), few studies have demonstrated efficacy in pre-clinical models. Rhesus macaques immunized with two or three doses of a DNA plasmid vaccine encoding full-length SARS-CoV-2 S protein induced neutralizing antibody in sera and reduced viral burden in BAL and nasal mucosa fluids (Yu et al., 2020). Moreover, three immunizations over two weeks with purified, inactivated SARS-CoV-2 induced neutralizing antibodies, and depending on the dose administered provided partial or complete protection against infection and viral pneumonia in rhesus macaques (Gao et al., 2020). One limitation of these challenge models is that rhesus macaques develop mild interstitial pneumonia after SARS-CoV-2 infection compared to some humans and other non-human primate species (Chandrashekar et al., 2020; Munster et al., 2020). Our study in hACE2-expressing mice show that a single intramuscular or intranasal dose of ChAd-SARS-CoV-2-S vaccine confers substantial and possibly complete protection against viral replication, inflammation, and lung disease.

Our approach supports the use of non-human Ad-vectored vaccines against emerging RNA viruses including SARS-CoV-2. Previously, we showed the efficacy of single-dose or two-dose regimens of a gorilla Ad encoding the prM-E genes of Zika virus (ZIKV) in several mouse challenge models including in the context of pregnancy (Hassan et al., 2019). Others have evaluated ChAd or rhesus macaque Ad vaccine candidates against ZIKV and shown efficacy in mice and non-human primates (Abbink et al., 2016; López-Camacho et al., 2018). A different ChAd encoding the wild-type SARS-CoV-2 S protein (ChAdOx1) is currently in clinical trials in humans (NCT04324606). Although data from the human trials has not yet been reported, studies in rhesus macaques suggest that a single intramuscular dose protects against infection in the lung but not in the upper respiratory tract (https://www.biorxiv.org/content/10.1101/2020.05.13.093195v1). None of the vaccines evaluated against SARS-CoV-2 has shown evidence of immune enhancement in any pre-clinical or clinical study, which has been a theoretical risk based on studies with other human and animal coronaviruses (de Alwis et al., 2020; Diamond and Pierson, 2020). Indeed, and in contrast to data with SARS-CoV vaccines or antibodies (Bolles et al., 2011; Liu et al., 2019; Weingartl et al., 2004), we did not observe enhanced infection, immunopathology, or disease in animals immunized with ChAd encoding SARS-CoV-2 S proteins or administered passively transferred monoclonal antibodies (Alsoussi et al., 2020; Hassan et al., 2020).

ChAd-SARS-CoV-2-S induced SARS-CoV-2 specific CD8^+^ T cell responses including a high percentage and number of IFNγ and granzyme expressing cells after *ex vivo* S protein peptide restimulation. The induction of robust CD8^+^ T cell responses by the ChAd-SARS-CoV-2-S vaccine is consistent with reports with other simian Ad vectors (Douglas et al., 2010; Hodgson et al., 2015; Reyes-Sandoval et al., 2010). ChAd vaccine vectors not only overcome issues of pre-existing immunity against human adenoviruses but also have immunological advantages because they do not induce exhausted T cell responses (Penaloza-MacMaster et al., 2013).

A single intranasal dose of ChAd-SARS-CoV-2-S conferred superior immunity against SARS-CoV-2 challenge, more so than one or two intramuscular immunizations of the same vaccine and dose. Given that the serum neutralizing antibody responses was comparable, we hypothesize the greater protection observed after intranasal delivery was because of the mucosal immune responses generated. Indeed, high levels of anti-SARS-CoV-2 IgA were detected in serum and lung, and B cells secreting IgA were detected in the spleen. Moreover, intranasal vaccination also induced SARS-CoV-2-specific CD8^+^ T cells in the lung including CD103^+^CD69^+^ cells, which are likely of a resident memory phenotype. To our knowledge, none of the SARS-CoV-2 vaccine platforms currently in clinical trials use an intranasal delivery approach. There has been great interest in using intranasal delivery for influenza A virus vaccines because of their ability to elicit local humoral and cellular immune responses (Calzas and Chevalier, 2019). Indeed, sterilizing immunity to influenza A virus re-infection requires local adaptive immune responses in the lung, which is optimally induced by intranasal and not intramuscular inoculation (Dutta et al., 2016; Laurie et al., 2010). While there are concerns with administering live-attenuated viral vaccines via an intranasal route, subunit-based or replication-incompetent vectored vaccines are promising for generating mucosal immunity in a safer manner, especially given advances in formulation (Yusuf and Kett, 2017).

Although a single intranasal administration of ChAd-SARS-CoV-2-S protected against SARS-CoV-2 replication in lungs, we note some limitations in our study. We performed challenge studies in mice transduced with hACE2 using a human Ad5 vector, which could be complicated by vector immunity. Notwithstanding this possibility, we confirmed an absence of heterologous neutralizing serum antibody induced by one or two doses of ChAd vector against the Hu-Ad5 and more importantly, detected high levels of SARS-CoV-2 infection in the ChAd-control vaccinated mice after challenge. Future immunization and challenge studies with hACE2 transgenic mice (Bao et al., 2020; Jiang et al., 2020) and non-human primates can overcome this potential issue and expand upon our understanding of how route of inoculation impacts vaccine-mediated protection. Moreover, studies in humans are needed to assess for cross-immunity between the ChAd vector (simian Ad-36) and adenoviruses circulating in humans. Although we expect this will be much less than that observed with Hu-Ad5 vaccine vectors (Zhu et al., 2020), low levels of pre-existing immunity against simian adenoviruses could affect vaccine efficacy in selected populations with non-human primate exposures (*e*.*g*., zoo workers, veterinarians, and laboratory workers at primate centers). Finally, studies must be conducted to monitor immune responses over time after intranasal vaccination with ChAd-SARS-CoV-2-S to establish the durability of the protective response.

In summary, our studies establish that immunization with ChAd-SARS-CoV-2-S induces both neutralizing antibody and antigen-specific CD8^+^ T cell responses. While a single intramuscular immunization of ChAd-SARS-CoV-2-S confers protection against SARS-CoV-2 infection and inflammation in the lungs, intranasal delivery of ChAd-SARS-CoV-2-S induces mucosal immunity, provides superior protection, and likely promotes sterilizing immunity, at least in mice transiently expressing the hACE2 receptor. We suggest that intranasal delivery of ChAd-SARS-CoV-2-S is a promising platform for controlling SARS-CoV-2 infection, disease, and transmission, and thus warrants clinical evaluation in humans.

## ACKNOWLEDGEMENTS

This study was supported by NIH contracts and grants (75N93019C00062, R01 AI127828, R01 AI130591, R01 AI149644, and R35 HL145242) and the Defense Advanced Research Project Agency (HR001117S0019). B.T.M is supported by F32 AI138392, E.S.W. is supported by T32 AI007163, and J.B.C. is supported by a Helen Hay Whitney Foundation postdoctoral fellowship. We thank Sean Whelan, Susan Cook, and Jennifer Philips for facilitating the high-containment studies with SARS-CoV-2, James Earnest for performing cell culture studies, and the Pulmonary Morphology Core at Washington University School of Medicine for tissue sectioning and slide preparation.

## AUTHOR CONTRIBUTIONS

A.O.H. amplified the adenovirus, designed experiments, and performed intranasal inoculations of adenovirus. I.P.D., E.K., and D.T.C designed and generated the ChAd constructs. J.B.C. propagated the SARS-CoV-2 stocks, developed the focus forming, neutralization, and plaque assays, and performed intranasal inoculations of SARS-CoV-2. A.L.B. designed and constructed the RT-qPCR assays. N.M.K., E.S.W., S.S., B.T.M., R.E.C., L.E.G., and J.M.F. performed clinical analysis, tissue harvests, histopathological studies, and viral burden analyses. B.T.M. performed *in situ* hybridization. A.S. and Y.J.H. designed and performed experiments with recombinant strains of SARS-CoV-2. A.S.K. performed flow cytometry analysis experiments. S.P.K. and M.J.H. analyzed the tissue sections for histopathology. A.W.W. performed the ChAd-immune serum neutralization assays against Hu-Ad5. D.W. generated the peptide pools for T cell restimulation. L.A.V.B. provided the anti-SARS-CoV-2 monoclonal antibodies. H.Z., Y.D., M.M., L.J.A., and D.H.F. designed and produced the recombinant S, RBD, and N proteins. I.B.H. and B.S. performed ELISAs, and J.B.C. and R.E.C. performed neutralization assays. R.B. designed experiments and secured funding. A.O.H. and M.S.D. wrote the initial draft, with the other authors providing editorial comments.

## COMPETING FINANCIAL INTERESTS

M.S.D. is a consultant for Inbios, Vir Biotechnology, NGM Biopharmaceuticals, and on the Scientific Advisory Board of Moderna. The Diamond laboratory has received unrelated funding support in sponsored research agreements from Moderna, Vir Biotechnology, and Emergent BioSolutions. M.J.H. is a member of the DSMB for AstroZeneca and founder of NuPeak Therapeutics. The Baric laboratory has received unrelated funding support in sponsored research agreements with Takeda, Pfizer, and Eli Lily.

## STAR METHODS

### RESOURCE AVAILABLITY

#### Lead Contact

Further information and requests for resources and reagents should be directed to and will be fulfilled by the Lead Contact, Michael S. Diamond.

#### Materials Availability

All requests for resources and reagents should be directed to and will be fulfilled by the Lead Contact author. This includes mice, antibodies, viruses, vaccines, proteins, and peptides. All reagents will be made available on request after completion of a Materials Transfer Agreement.

#### Data and code availability

All data supporting the findings of this study are available within the paper and are available from the corresponding author upon request.

## EXPERIMENTAL MODEL AND SUBJECT DETAILS

### Viruses and cells

Vero E6 (CRL-1586, American Type Culture Collection (ATCC), Vero CCL81 (ATCC), and HEK293 cells were cultured at 37°C in Dulbecco’s Modified Eagle medium (DMEM) supplemented with 10% fetal bovine serum (FBS), 10 mM HEPES pH 7.3, 1 mM sodium pyruvate, 1X non-essential amino acids, and 100 U/ml of penicillin–streptomycin.

SARS-CoV-2 strain 2019 n-CoV/USA_WA1/2020 was obtained from the Centers for Disease Control and Prevention (a gift from Natalie Thornburg). The virus was passaged once in Vero CCL81 cells and titrated by focus-forming assay (FFA) on Vero E6 cells. The recombinant luciferase-expressing full-length SARS-CoV-2 reporter virus (2019 n-CoV/USA_WA1/2020 strain) has been reported previously (Zost et al., 2020), and the D614G variant will be described elsewhere (R. Baric, manuscript in preparation). All work with infectious SARS-CoV-2 was performed in Institutional Biosafety Committee approved BSL3 and A-BSL3 facilities using appropriate positive pressure air respirators and protective equipment.

### Mouse experiments

Animal studies were carried out in accordance with the recommendations in the Guide for the Care and Use of Laboratory Animals of the National Institutes of Health. The protocols were approved by the Institutional Animal Care and Use Committee at the Washington University School of Medicine (Assurance number A3381-01). Virus inoculations were performed under anesthesia that was induced and maintained with ketamine hydrochloride and xylazine, and all efforts were made to minimize animal suffering.

Female BALB/c mice were purchased from The Jackson Laboratory (catalog 000651). Four to five-week-old animals were immunized with 10^10^ viral particles (vp) of ChAdV-empty or ChAd-SARS-CoV-2-S in 50 µl PBS via intramuscular injection in the hind leg or via intranasal inoculation. Subsets of immunized animals were boosted four weeks after primary immunization using the same route used for primary immunization. Vaccinated mice (10 to 11-week-old) were given a single intraperitoneal injection of 2 mg of anti-Ifnar1 mAb (MAR1-5A3 (Sheehan et al., 2006), Leinco) one day before intranasal administration of 10^8^ PFU of Hu-AdV5-hACE2. Five days after Hu-AdV5 transduction, mice were inoculated with 4 × 10^5^ FFU of SARS-CoV-2 by the intranasal route. Animals were euthanized at 4 or 8 dpi, and tissues were harvested for virological, immunological, and pathological analyses.

## METHOD DETAILS

### Construction of chimpanzee adenovirus vectors

Simian Ad36 vector (ChAd) (Roy et al., 2011) was obtained from the Penn Vector Core of the University of Pennsylvania. The ChAd genome was engineered with deletions in the E1 and E3B region (GenBank: FJ025917.1; nucleotides 455-3026 and 30072-31869, respectively). A modified human cytomegalovirus major immediate early promoter sequence was incorporated in place of the E1 gene in counterclockwise orientation on the complementary DNA strand. CMV modification included an addition of two copies of the tet operator 2 (TetO2) sequence (Hillen and Berens, 1994) inserted in tandem (5’-TCT CTA TCA CTG ATA GGG AGA TCT CTA TCA CTG ATA GG GA-3’) between the TATA box and the mRNA start (GenBank: MN920393, nucleotides 174211-174212). SARS-CoV-2 S (encoding a prefusion stabilized mutant with two proline substitutions at residues K986 and V987 that stabilize the prefusion form of S (Pallesen et al., 2017; Wrapp et al., 2020a)) was cloned into a unique *Pme*I site under the CMV-tetO2 promoter control in pSAd36 genomic plasmid to generate pSAd36-S. In parallel, a pSAd36-control carrying an empty CMV-tetO2 cassette with no transgene also was generated. The pSAd36-S and pSAd-control plasmids were linearized with *Pac*I restriction enzyme to liberate viral genomes for transfection into T-Rex 293-HEK cells (Invitrogen). The rescued replication-incompetent ChAd-SARS-CoV-2-S and ChAd-Control vectors were scaled up in 293 cells and purified by CsCl density-gradient ultracentrifugation. Viral particle concentration in each vector preparation was determined by spectrophotometry at 260 nm as described (Maizel et al., 1968).

### Construction of a human adenovirus vector expressing human ACE2

Codon-optimized hACE2 sequences were cloned into the shuttle vector (pShuttle-CMV, Addgene 240007) to generate pShuttle-hACE2. pShuttle-hACE2 was linearized with *Pme*I and subsequently cotransformed with the HuAdv5 backbone plasmid (pAdEasy-1 vector; Addgene 240005) into *E. coli* strain BJ5183 to generate pAdV5-ACE2 by homologous recombination as described (He et al., 1998). The pAdEasy-1 plasmid containing the HuAdV5 genome has deletions in E1 and E3 genes. hACE2 is under transcriptional control of a cytomegalovirus promotor and is flanked at its 3’ end by a SV40 polyadenylation signal. The pAd-hACE2 was linearized with *Pac*I restriction enzyme before transfection into T-Rex 293 HEK cells (Invitrogen) to generate HuAdv5-hACE2. Recombinant HuAdv5-hACE2 was produced in 293-HEK cells and purified by CsCl density-gradient ultracentrifugation. The viral titer was determined by plaque assay in 293-HEK cells.

### In situ RNA hybridization and histology

RNA *in situ* hybridization was performed using RNAscope 2.5 HD (Brown) (Advanced Cell Diagnostics) according to the manufacturer’s instructions. Left lung tissues were collected at necropsy, inflated with 10% neutral buffered formalin (NBF), and thereafter immersion fixed in 10% NBF for seven days before processing. Paraffin-embedded lung sections were deparaffinized by incubating at 60°C for 1 h, and endogenous peroxidases were quenched with H_2_O_2_ for 10 min at room temperature. Slides were boiled for 15 min in RNAscope Target Retrieval Reagents and incubated for 30 min in RNAscope Protease Plus reagent prior to SARS-CoV2 RNA probe (Advanced Cell Diagnostics 848561) hybridization and signal amplification. Sections were counterstained with Gill’s hematoxylin and visualized by brightfield microscopy. Some lung sections were processed for histology after hematoxylin and eosin staining.

### SARS-CoV-2 Neutralization assays

Heat-inactivated serum samples were diluted serially and incubated with 10^2^ FFU of SARS-CoV-2 for 1 h at 37°C. The virus-serum mixtures were added to Vero cell monolayers in 96-well plates and incubated for 1 h at 37°C. Subsequently, cells were overlaid with 1% (w/v) methylcellulose in MEM supplemented with 2% FBS. Plates were incubated for 30 h before fixation using 4% PFA in PBS for 1 h at room temperature. Cells were washed then sequentially incubated with anti-SARS-CoV-2 CR3022 antibody (Yuan et al., 2020) (1 μg/mL) and a HRP-conjugated goat anti-human IgG (Sigma) in PBS supplemented with 0.1% (w/v) saponin (Sigma) and 0.1% BSA. TrueBlue peroxidase substrate (KPL) was used to develop the plates before counting the foci on a BioSpot analyzer (Cellular Technology Limited). For neutralization experiments with luciferase expressing SARS-CoV-2, serum samples were diluted 3-fold starting at 1:50 and mixed with 85 PFU of each recombinant virus (wild-type and D614G). Vero E6 cells plated in clear-bottom black-walled 96-well plates (Corning) were inoculated with serum-virus mixtures, and cells were cultured at 37°C for 48 h. Subsequently, cells were lysed and luciferase activity was measured using the Nano-Glo Luciferase Assay System (Promega) according to the manufacturer’s specifications.

### Hu-AdV5 neutralization assays

One day prior to Hu-AdV5-hACE2 transduction, serum samples were collected from mice immunized intramuscularly with ChAd-Control or ChAd-SARS-CoV-2-S. Sera were heat-inactivated and serially diluted prior to incubation with 10^2^ FFU of Hu-AdV5 for 1 h at 37°C. The virus-serum mixtures were added to HEK293 cell monolayers in 96-well plates and incubated for 1 h at 37°C. Cells were then overlaid with 1% (w/v) methylcellulose in MEM supplemented with 5% FBS. Plates were incubated at 37°C for 48 h before fixation with 2% PFA in PBS for 1 h at room temperature. Subsequently, plates were washed with PBS and incubated overnight at 4°C with biotinylated anti-HuAdV5-hexon antibody (2 µg/mL; Novus Biologicals NB600413) diluted in permeabilization buffer (PBS supplemented with 0.1% (w/v) saponin and 0.1% BSA). Plates were washed again and incubated with streptavidin-HRP (1:3000; Vector Laboratories SA-5004) in permeabilization buffer for 30 min at room temperature. After a final wash series, plates were developed using TrueBlue peroxidase substrate (KPL) and foci were counted on a BioSpot analyzer (Cellular Technology Limited).

### Protein expression and purification

Purified RNA from the 2019-nCoV/USA-WA1/2020 SARS-CoV-2 strain was reverse transcribed into cDNA and used as the template for recombinant gene cloning. A full-length SARS-CoV-2 NP (NP-FL) was cloned into pET21a with a hexahistidine tag and recombinantly expressed using BL21(DE3)-RIL *E. coli* in Terrific Broth (bioWORLD). Following overnight induction with isopropyl β-d-1-thiogalactopyranoside (Goldbio) at 25°C, cells were lysed in 20 mM Tris-HCl pH 8.5, 1 M NaCl, 5 mM β-mercaptopethanol, and 5 mM imidazole for nickel-affinity purification. Following elution in the prior buffer supplemented with 500 mM imidazole, the protein was purified to homogeneity using size exclusion and, in some cases, cation exchange chromatography. SARS-CoV-2 RBD and S ectodomain (S1/S2 furin cleavage site was disrupted, double proline mutations were introduced into the S2 subunit, and foldon trimerization motif was incorporated) were cloned into pFM1.2 with a C-terminal hexahistidine or octahistidine tag, transiently transfected into Expi293F cells, and purified by cobalt-charged resin chromatography (G-Biosciences) as previously described (Alsoussi et al., 2020).

### ELISA

Purified antigens (S, RBD, or NP) were coated onto 96-well Maxisorp clear plates at 2 µg/mL in 50 mM Na_2_CO_3_pH 9.6 (70 µL) overnight at 4°C. Coating buffers were aspirated, and wells were blocked with 200 µL of 1X PBS + 0.05% Tween-20 + 1% BSA + 0.02% NaN_3_ (Blocking buffer, PBSTBA) either for 1 h at 37°C or overnight at 4°C. Heat-inactivated serum samples were diluted in PBSTBA in a separate 96-well polypropylene plate. The plates then were washed thrice with 1X PBS + 0.05% Tween-20 (PBST), followed by addition of 50 µL of respective serum dilutions. Sera were incubated in the blocked ELISA plates for at least 1 h at room temperature. The ELISA plates were again washed thrice in PBST, followed by addition of 50 µL of 1:2000 anti-mouse IgG-HRP (Southern Biotech Cat. #1030-05) in PBST or 1:10000 of biotinylated anti-mouse IgG, anti-mouse IgM, or anti-mouse IgA in PBSTBA (SouthernBiotech). Plates were incubated at room temperature for 1 h, washed thrice in PBST, and then 1:5000 dilution of streptavidin-HRP (ThermoFisher) was added to wells. Following a 1 h incubation at room temperature, plates were washed thrice with PBST and 50 µL of 1-Step Ultra TMB-ELISA was added (ThermoFisher Cat. #34028). Following a 12 to 15-min incubation, reactions were stopped with 50 µL of 2 M sulfuric acid. The absorbance of each well at 450 nm was read (Synergy H1) within 2 min of addition of sulfuric acid. Optical density (450 nm) measurements were determined using a microplate reader (Bio-Rad).

### ELISpot assay

MultiScreen-HA filter 96-well plates (Millipore) plates were pre-coated with 3 µg/ml of SARS-CoV-2 S protein overnight at 4°C. After rinsing with PBST, plates were blocked for 4 h at 37°C with culture medium (RPMI, 10% FBS, penicillin-streptomycin, 1 mM sodium pyruvate, 0.1 mM non-essential amino acids, 10 mM HEPES, and 50 mM β-mercaptoethanol). Single cell suspensions of splenocytes in culture medium were added to the S protein coated plates and incubated at 37°C and 5% humidified CO_2_ for 4 h. After washing with PBS and PBST, plates were incubated with biotinylated anti-IgG or anti-IgA (Southern Biotech) followed by incubation with streptavidin conjugated horseradish peroxidase (Jackson ImmunoResearch), each for 1 h at room temperature. After additional washes with PBS, 3-amino-9-ethylcarbazole (Sigma) substrate solution was added for spot development. The reaction was stopped by rinsing with water. Spots were counted using a Biospot plate reader (Cellular Technology).

### Measurement of viral burden

SARS-CoV-2 infected mice were euthanized using a ketamine and xylazine cocktail, and organs were collected. Tissues were weighed and homogenized with beads using a MAgNA Lyser (Roche) in 1 ml of Dulbecco’s Modified Eagle’s Medium (DMEM) containing 2% fetal bovine serum (FBS). RNA was extracted from clarified tissue homogenates using MagMax mirVana Total RNA isolation kit (Thermo Scientific) and a Kingfisher duo prime extraction machine (Thermo Scientific). SARS-CoV-2 RNA levels were measured by one-step quantitative reverse transcriptase PCR (qRT-PCR) TaqMan assay as described previously (Hassan et al., 2020). SARS-CoV-2 nucleocapsid (N) specific primers and probe set were used: (L Primer: ATGCTGCAATCGTGCTACAA; R primer: GACTGCCGCCTCTGCTC; probe: /56-FAM/TCAAGGAAC/ZEN/AACATTGCCAA/3IABkFQ/) (Integrated DNA Technologies). Viral RNA was expressed as (N) gene copy numbers per milligram on a log_10_ scale. For some samples, viral titer was determined by plaque assay on Vero E6 cells.

### Cytokine and chemokine mRNA measurements

RNA extracted from lung homogenates lung homogenates was DNAse-treated and used to synthesize cDNA using the High-Capacity cDNA Reverse Transcription kit (Thermo Scientific) with the addition of RNase inhibitor according to the manufacturer’s protocol. Cytokine and chemokine expression was determined using TaqMan Fast Universal PCR master mix (Thermo Scientific) with commercial primers/probe sets specific for *IFN-γ* (IDT: Mm.PT.58.41769240), *IL-6* (Mm.PT.58.10005566), *IL-1β* (Mm.PT.58.41616450), *TNF-α* (Mm.PT.58.12575861), *CXCL10* (Mm.PT.58.43575827), *CCL2* (Mm.PT.58.42151692), *CCL5* (Mm.PT.58.43548565), *CXCL11* (Mm.PT.58.10773148.g), *IFN-β* (Mm.PT.58.30132453.g), and *IFNλ*−*2/3* (Thermo Scientific Mm04204156_gH) and results were normalized to *GAPDH* (Mm.PT.39a.1) levels. Fold change was determined using the 2^-ΔΔCt^ method comparing treated mice to naïve controls.

### Peptide restimulation and intracellular cytokine staining

Splenocytes from intramuscularly vaccinated mice were incubated in culture with a pool of 253 overlapping 15-mer SARS-CoV-2 S peptides (Grifoni et al., 2020) for 12 h at 37°C before a 4 h treatment with brefeldin A (BioLegend, 420601). Following blocking with FcγR antibody (BioLegend, clone 93), cells were stained on ice with CD45 BUV395 (BD BioSciences clone 30-F11); CD44 PE-Cy7, CD4 PE-Cy5, CD8b PreCP-Cy5.5, and CD19 APC-Cy7 (BioLegend clones, IM7, GK1.5, YTS156.7.7, and 6D5, respectively), and Fixable Aqua Dead Cell Stain (Invitrogen, L34966). Stained cells were fixed and permeabilized with the Foxp3/Transcription Factor Staining Buffer Set (eBiosciences, 00-5523). Subsequently, intracellular staining was performed with anti-IFN-γ Alexa 647 (BD Biosciences, clone XMG1.2), anti-TNFα BV605 (BioLegend, clone MP6-XT22), and anti-GrB PE (Invitrogen, GRB04). Lungs from intranasally immunized mice were harvested and digested for 1 h at 37°C in digestion buffer consisting of RPMI media supplemented with (167 μg/ml) of Liberase DH (Sigma) and (100 μg/ml) of DNase I (Sigma). Lung cells were incubated at 37°C with the pool of 253 overlapping 15-mer SARS-CoV-2 S peptides described above in the presence of brefeldin A for 5 h at 37°C. Lung cells then were stained as described above except that CD4-BV421 (BioLegend clone GK1.5) replaced the CD4-PE-Cy5, no CD19 staining was included, and CD103-FITC and CD69-BV711 (BioLegend clones, 2E7, and, H1.2F3, respectively) were added. Analysis was performed on a BD LSRFortessa X-20 cytometer, using FlowJo X 10.0 software.

### Flow cytometry-based antigen characterization

HEK-293T cells were seeded at 10^6^ cells/well in 6-well plates 24 h prior to transduction with ChAd-SARS-CoV-2-S (MOI of 5). After 20 h, cells were harvested, fixed and permeabilized using Foxp3 Transcription Factor Staining Buffer Set (Thermo Fisher), and stained for viral antigen after incubation with the following anti-SARS-CoV-2 neutralizing murine mAbs: SARS2-01, SARS2-02, SARS2-07, SARS2-11, SARS2-12, SARS2-16, SARS2-18, SARS2-20, SARS2-21, SARS2-22, SARS2-23, SARS2-29, SARS2-31, SARS2-32, SARS2-34, SARS2-38, SARS2-39, SARS2-50, SARS2-55, SARS2-58, SARS2-66, and SARS2-71 (L. VanBlargan and M. Diamond, unpublished results). H77.39 (Sabo et al., 2011), an isotype-matched anti-HCV E2 mAb was used as a negative control. Cells were washed, incubated with Alexa Fluor 647 conjugated goat anti-mouse IgG (Thermo Fisher), and analyzed by flow cytometry using a MACSQuant Analyzer 10 (Miltenyi Biotec). The percentage of cells positive for a given mAb was compared with cells stained with a saturating amount of an oligoclonal mixture of anti-SARS-CoV-2 mAbs.

## QUANTIFICATION AND STATISTICAL ANALYSIS

Statistical significance was assigned when *P* values were < 0.05 using Prism Version 8 (GraphPad). All tests and values are indicated in the relevant Figure legends.

## SUPPLEMENTARY FIGURE LEGENDS

**Figure S1.**
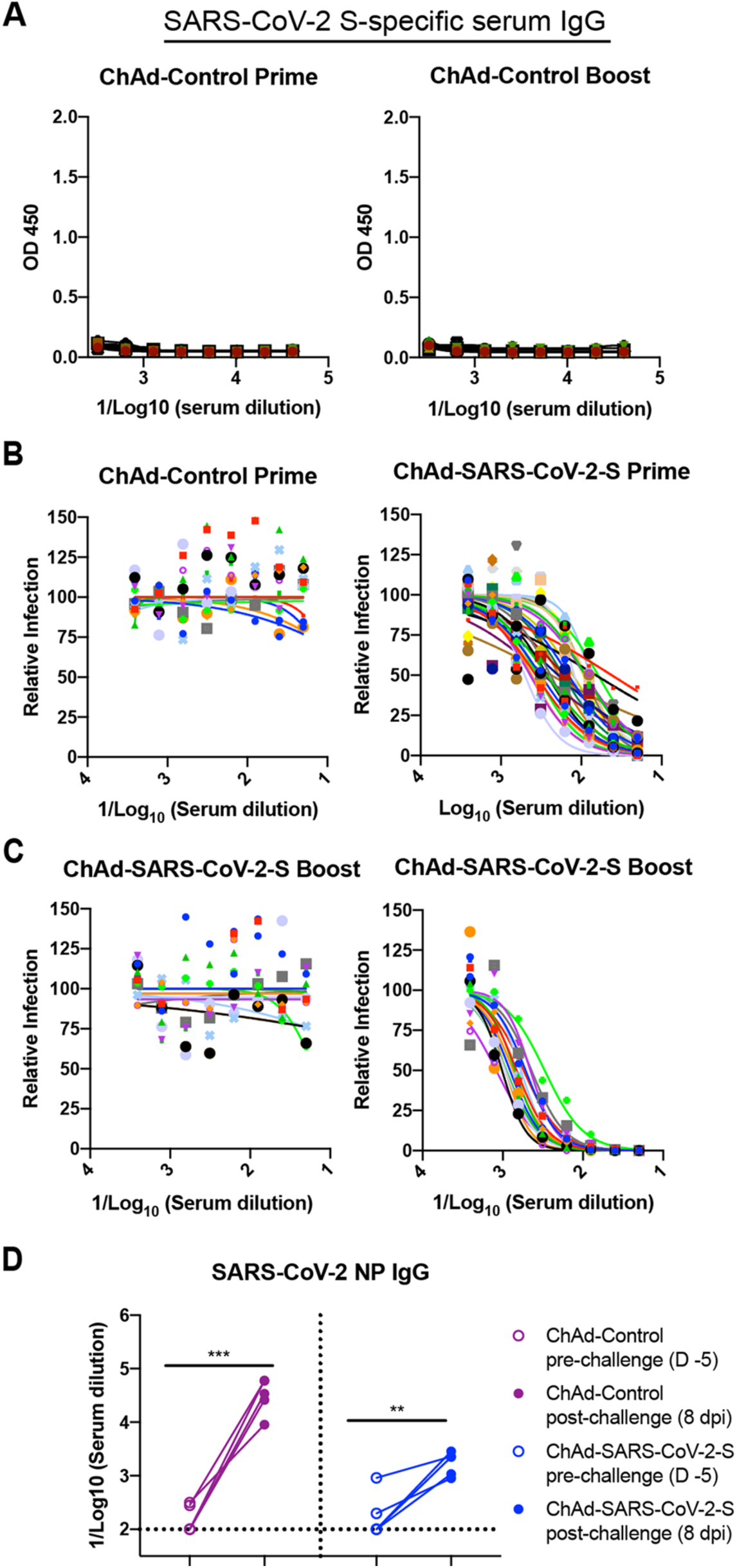
ChAd-SARS-CoV-2-S vaccine induces neutralizing antibodies as measured by FRNT. Related to Figure 1. Four-week old female BALB/c mice were primed or primed and boosted with ChAd-control or ChAd-SARS-CoV-2-S via intramuscular route. **A**. Serum from ChAd-control immunized mice collected at day 21 after priming or boosting (as described in **Fig 1**) was assayed for S-specific IgG responses by ELISA. **B-C**. Serum samples from ChAd-control or ChAd-SARS-CoV-2 vaccinated mice were collected at day 21 after priming (**B**) or boosting (**C**) and assayed for neutralizing activity by FRNT. Serum neutralization curves corresponding to individual mice are shown for the indicated vaccines (n = 15-30 per group). Each point represents the mean of two technical replicates with error bars denoting the standard deviations (SD). **D**. An ELISA measured anti-SARS-CoV-2 NP IgG responses in paired sera obtained 5 days before and 8 days after SARS-CoV-2 challenge of ChAd-control or ChAd-SARS-CoV-2-S mice vaccinated by an intramuscular route (n = 5: ** *P* < 0.01; *** *P* < 0.001; paired t test). Dotted lines represent the mean IgG titers from naïve sera.

**Figure S2.**
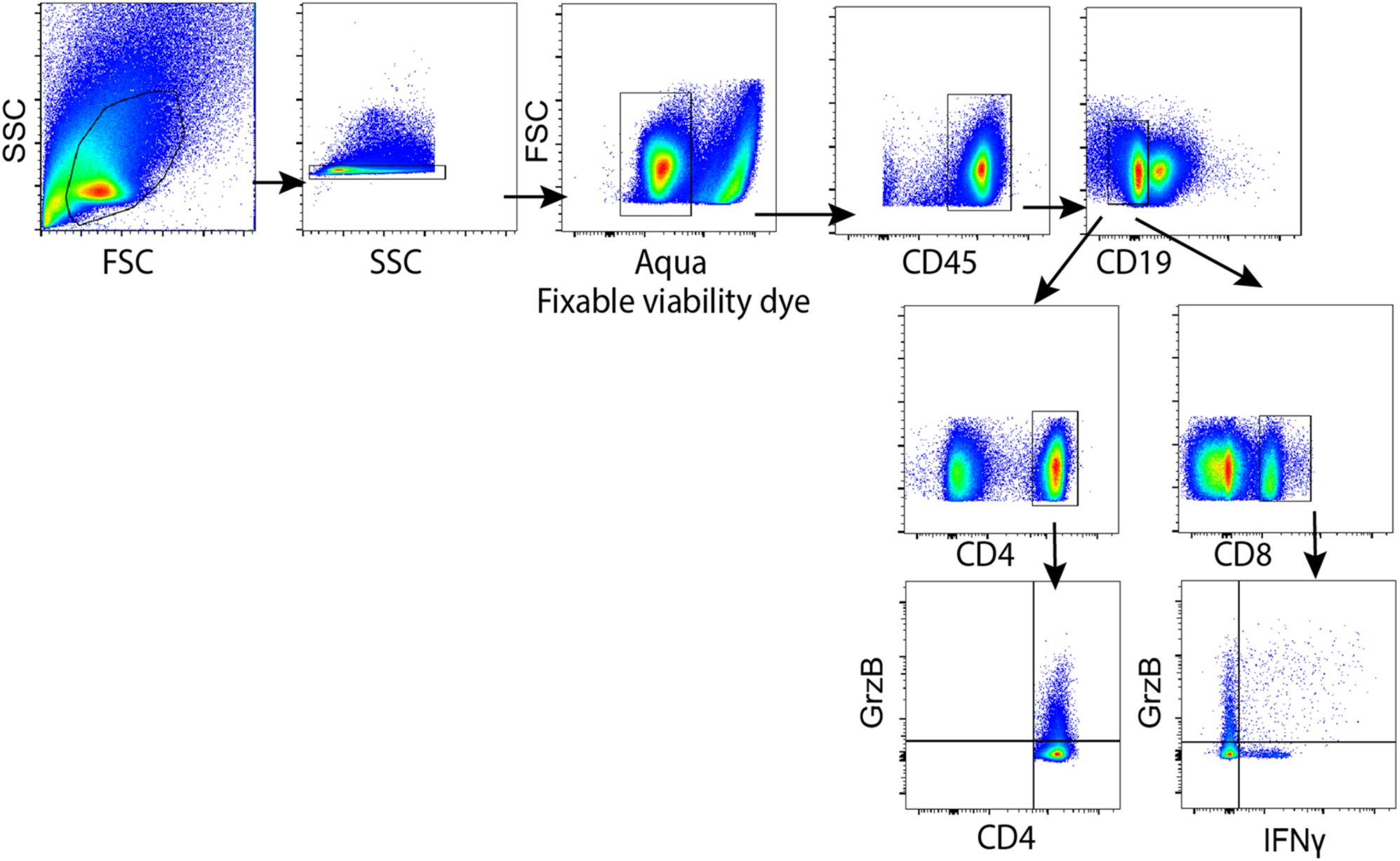
Gating strategy for analyzing T cell responses. Related to Figure 1. Four- week old female BALB/c mice were immunized with ChAd-control or ChAd-SARS-CoV-2-S and boosted four weeks later. T cell responses were analyzed in splenocytes at day 7 post-boost. Cells were gated for lymphocytes (FSC-A/SSC-A), singlets (SSC-W/SSC-H), live cells (Aqua^-^), CD45^+^, CD19^-^ followed by CD4^+^ or CD8^+^ cell populations expressing IFNγ or granzyme B.

**Figure S3.**
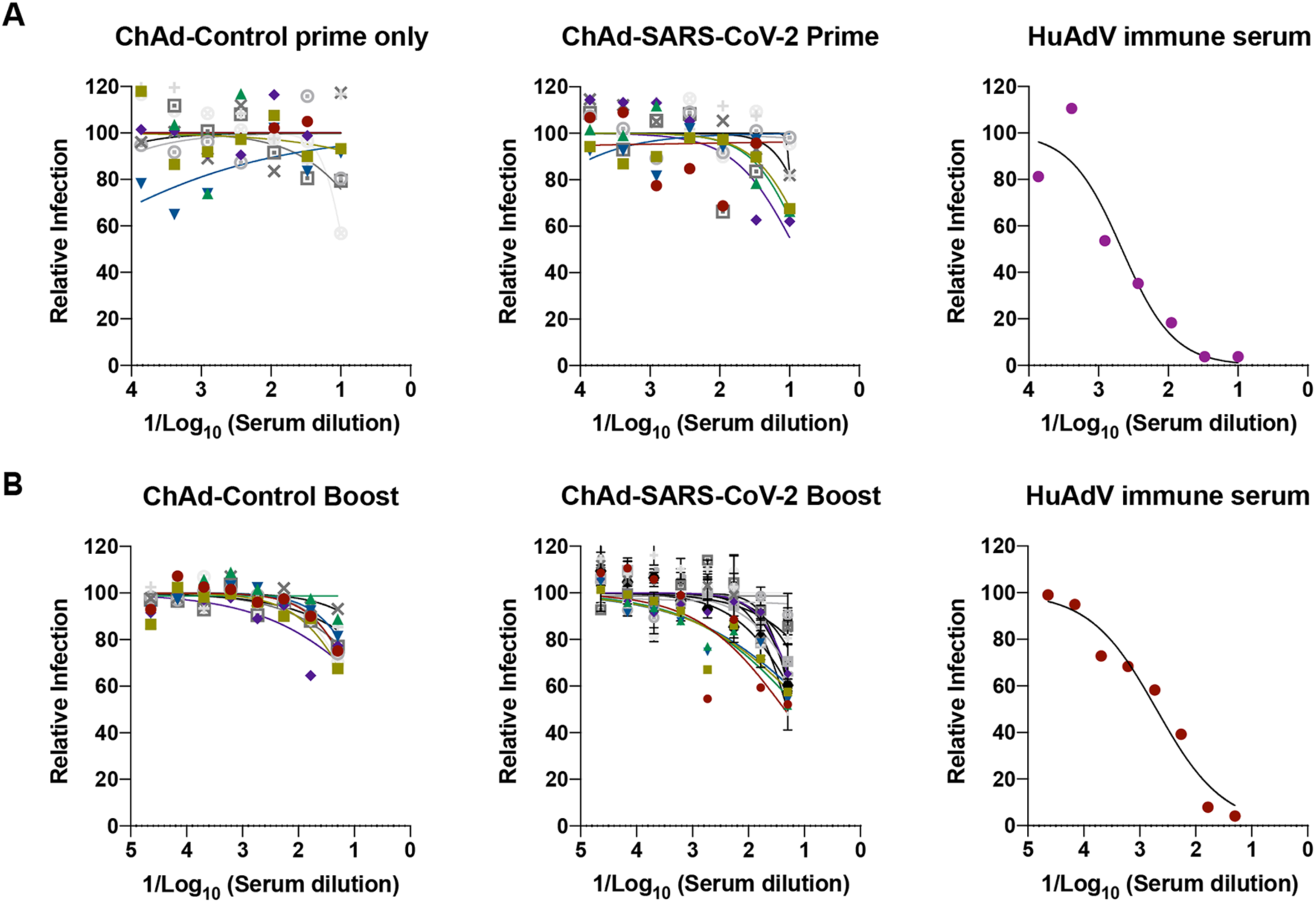
Impact of pre-existing ChAd immunity on transduction of mice with Hu-AdV5-hACE2. Related to Figures 1 and 2. Four-week old female BALB/c mice were primed or primed and boosted. Serum samples were collected one day prior to Hu-AdV5-hACE2 transduction. Neutralizing activity of Hu-AdV5-hACE2 in the sera from the indicated vaccine groups was determined by FRNT after prime only (**A**) or prime and boost (**B**). Each symbol represents a single animal; each point represents two technical repeats and bars indicate the range. A positive control (anti-Hu-Adv5 serum) is included as a frame of reference.

**Figure S4.**
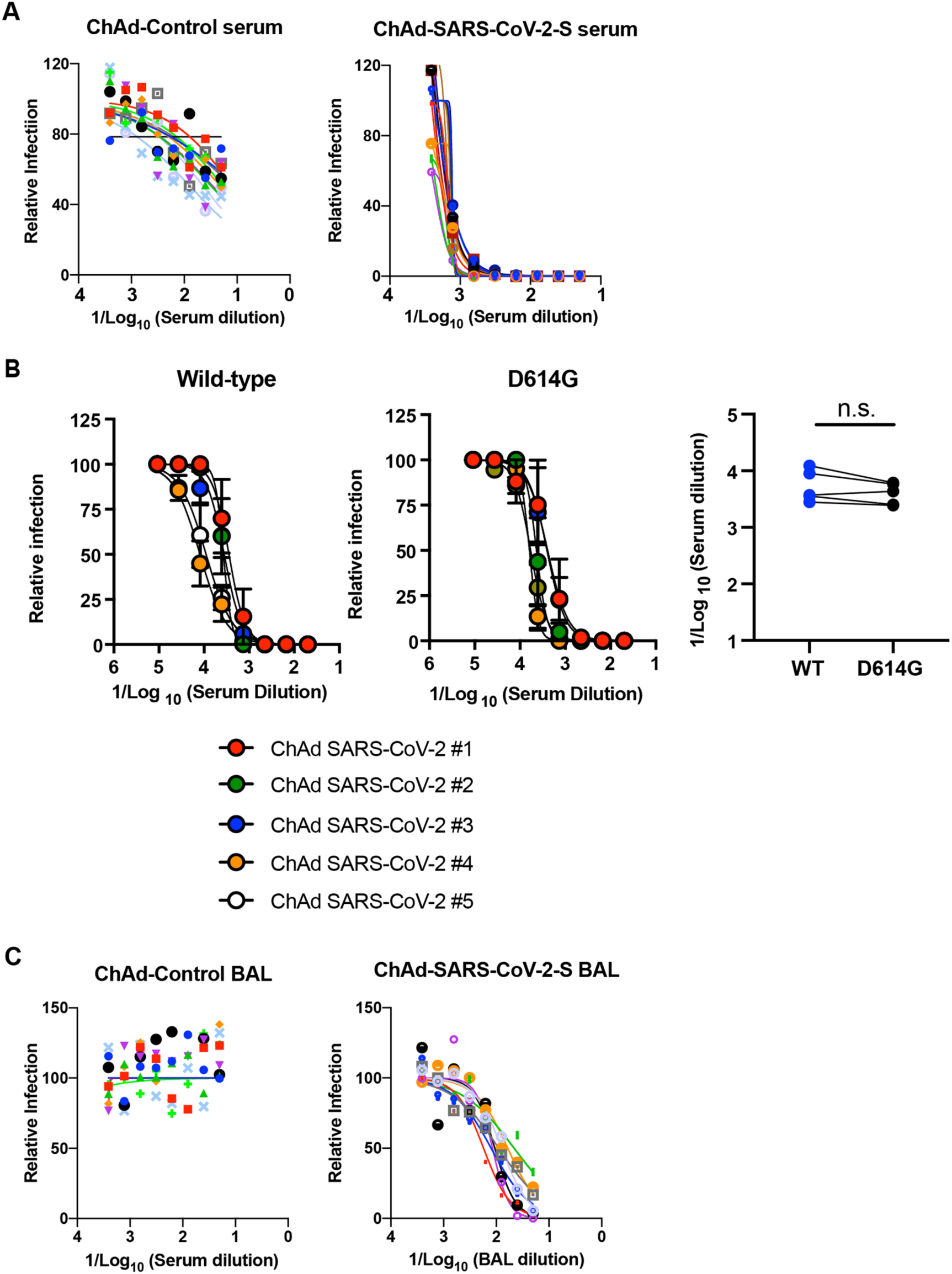
Intranasal inoculation of ChAd-SARS-CoV-2-S induces neutralizing antibodies as measured by FRNT. Related to Figure 4. Five-week old female BALB/c mice were immunized with ChAd-control or ChAd-SARS-CoV-2-S via an intranasal inoculation route. Serum samples collected one month after immunization were assayed for neutralizing activity by FRNT. Mice were boosted at day 30 after priming and were sacrificed one week later to evaluate immune responses. (**A**) Serum samples from ChAd-control or ChAd-SARS-CoV-2-S vaccinated mice were tested for neutralizing activity with SARS-CoV-2 strain 2019 n-CoV/USA_WA1/2020 (n = 8-10 per group). (**B**) Serum samples from ChAd-SARS-CoV-2-S vaccinated mice were tested for neutralization of recombinant luciferase-expressing SARS-CoV-2 viruses (wild-type (*left*) and D614G variant (*middle*)). (*Right*) Paired EC50 values are indicated (n = 5; n.s. not significant, paired t test). (**C**) BAL fluid was collected from ChAd-control or ChAd-SARS-CoV-2-S vaccinated mice, and neutralization of SARS-CoV-2 strain 2019 n-CoV/USA_WA1/2020 was measured using a FRNT assay (n = 8-10 per group). Each point represents the mean of two technical replicates with error bars denoting the SD.

**Table S1.**
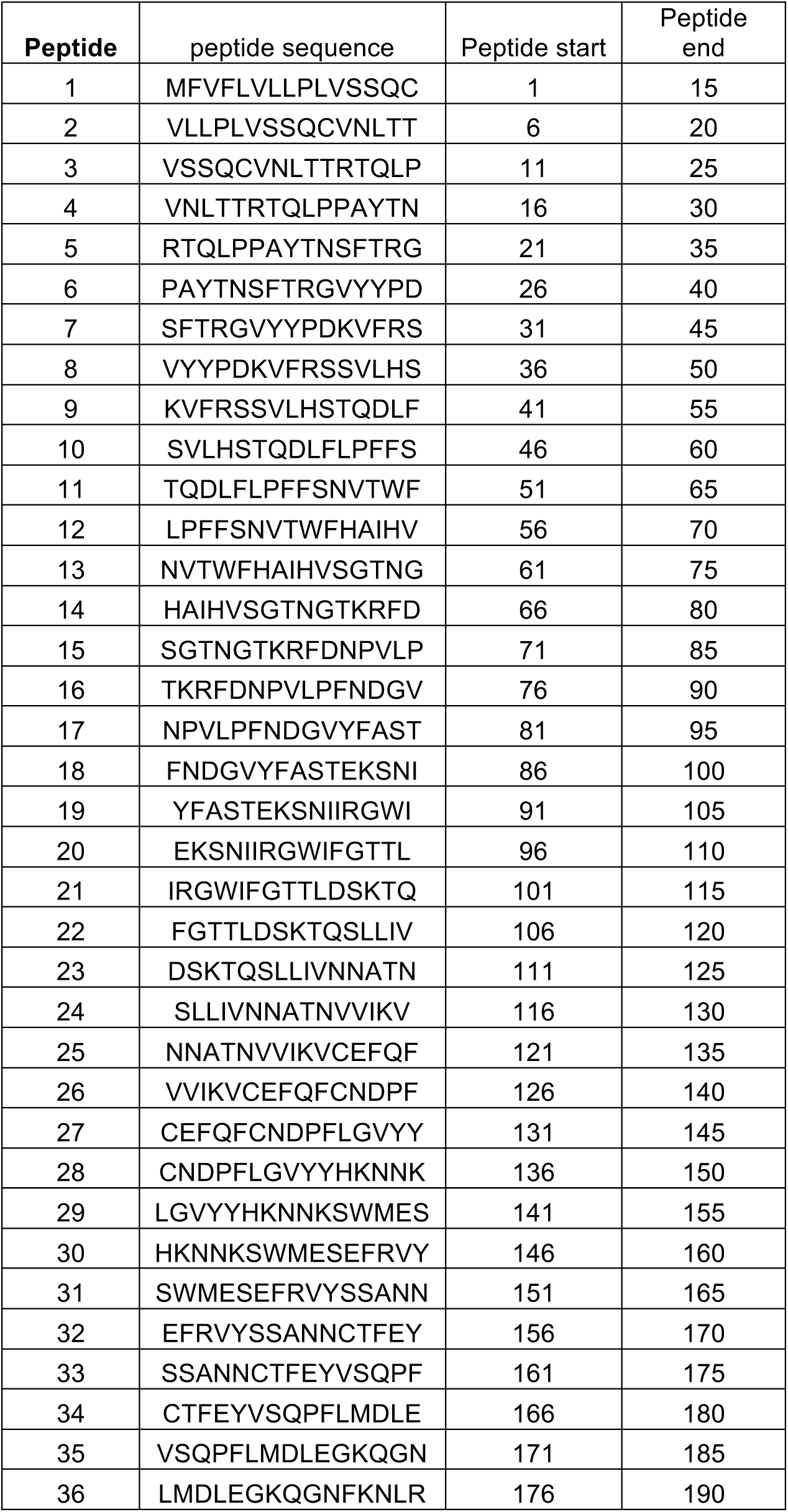
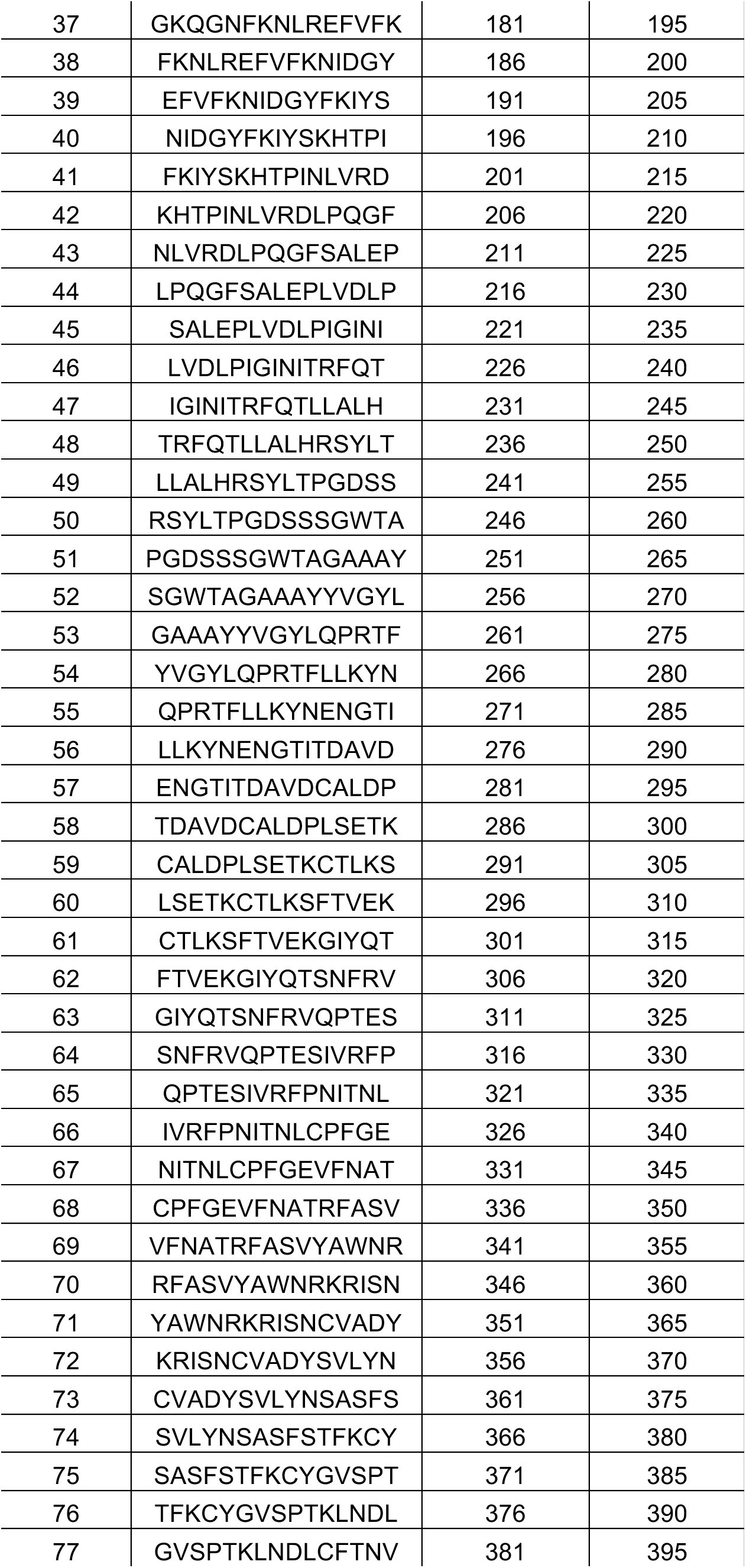
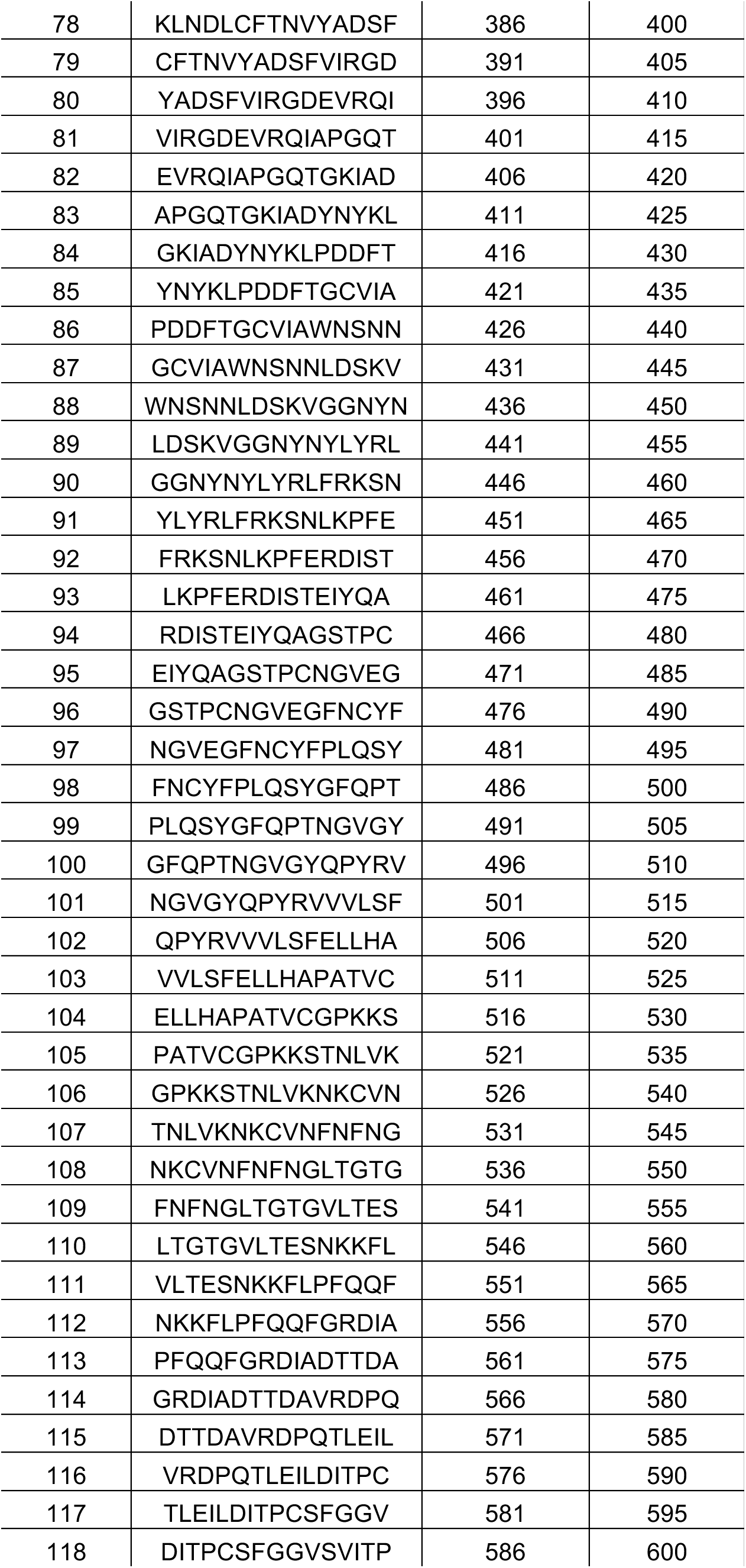
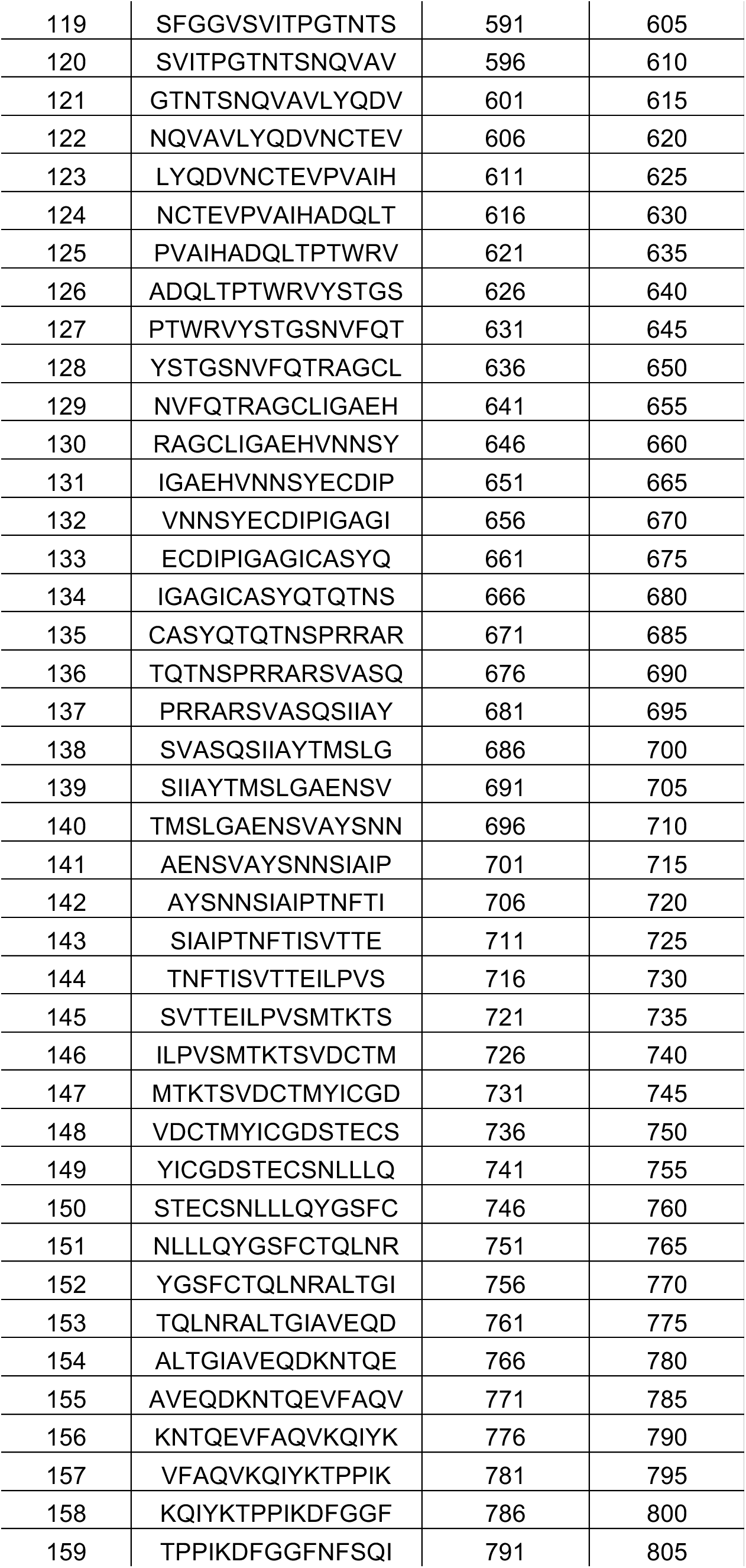
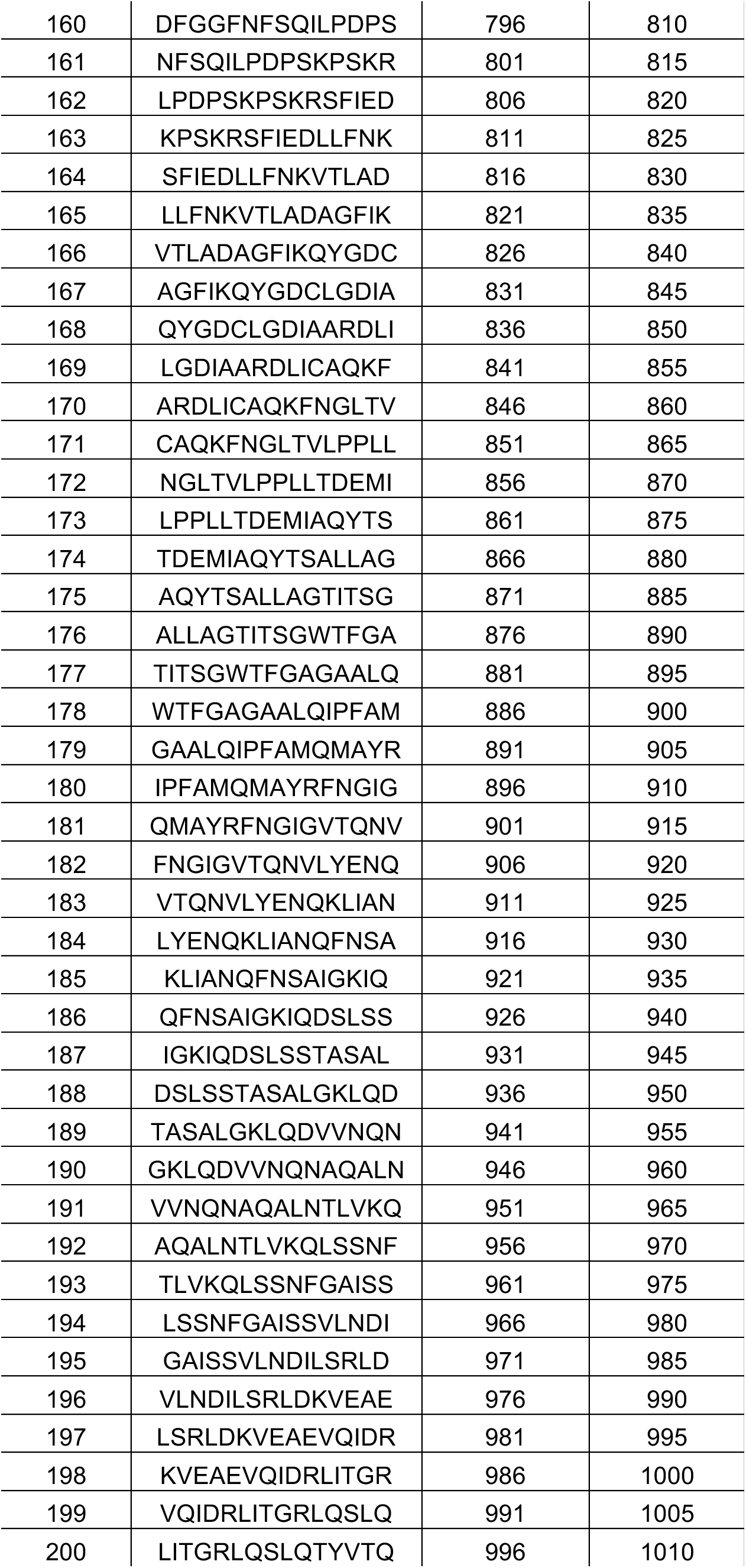
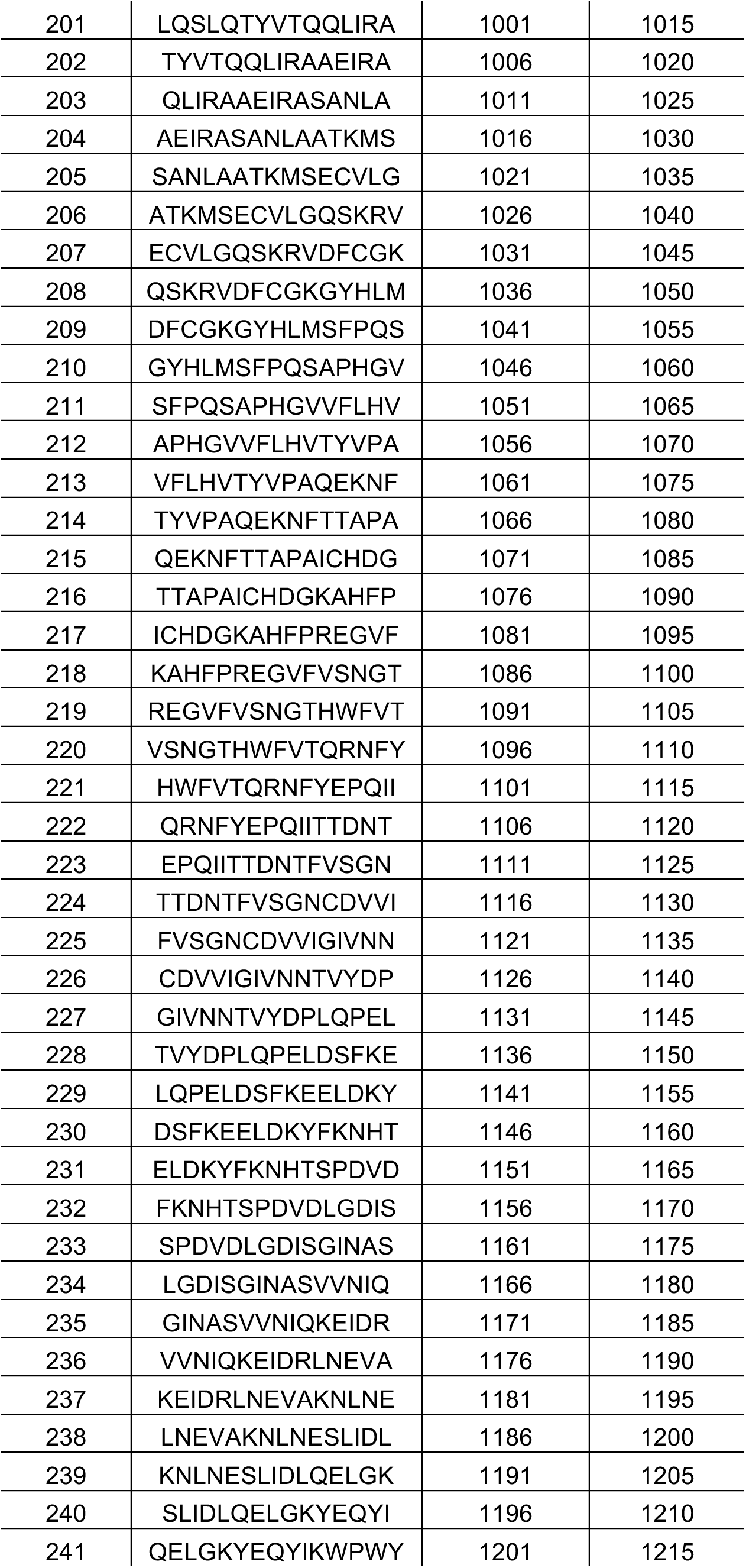
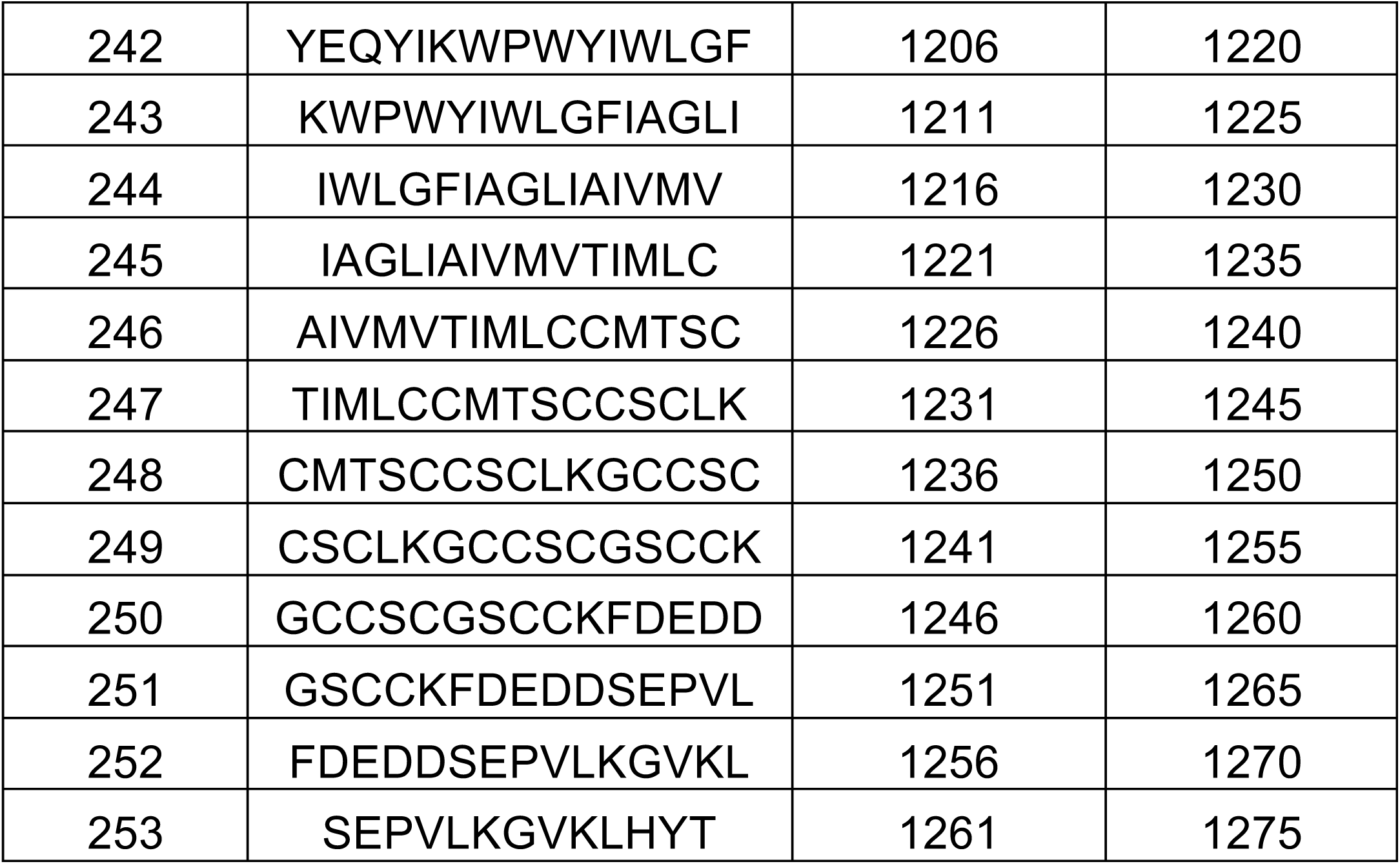
SARS-CoV-2 15-mer peptides in S protein

